# Accurate and Rapid Detection of Peritoneal Metastasis from Gastric Cancer by AI-assisted Stimulated Raman Cytology

**DOI:** 10.1101/2023.01.05.522829

**Authors:** Xun Chen, Zhouqiao Wu, Yexuan He, Zhe Hao, Qi Wang, Keji Zhou, Wanhui Zhou, Pu Wang, Fei Shan, Zhongwu Li, Jiafu Ji, Yubo Fan, Ziyu Li, Shuhua Yue

## Abstract

Peritoneal metastasis (PM) is the most common form of distant metastasis and one of the leading causes of death in gastric cancer (GC). For locally advanced GC, clinical guidelines recommend peritoneal lavage cytology for intraoperative PM detection. Unfortunately, current peritoneal lavage cytology is limited by low sensitivity (<60%). Here we established the stimulated Raman cytology (SRC), a chemical microscopy-based intelligent cytology. By taking advantages of stimulated Raman scattering in label-free, high-speed, and high-resolution chemical imaging, we firstly imaged 53951 exfoliated cells in ascites obtained from 80 GC patients (27 PM positive, 53 PM negative), at the Raman bands corresponding to DNA, protein, and lipid, respectively. Then, we revealed 12 single cell features of morphology and composition that were significantly different between PM positive and negative specimens, including cellular area, lipid protein ratio, etc. Importantly, we developed a single cell phenotyping algorithm to further transform the above raw features to feature matrix. Such matrix was crucial to identify the significant marker cell cluster, the divergence of which was finally used to differentiate the PM positive and negative. Compared with histopathology, the gold standard of PM detection, our SRC method assisted by machine learning classifiers could reach 81.5% sensitivity, 84.9% specificity, and the area under receiver operating characteristic curve of 0.85, within 20 minutes for each patient. Such remarkable improvement in detection accuracy is largely owing to incorporation of the single-cell composition features in SRC. Together, our SRC method shows great potential for accurate and rapid detection of PM from GC.

## Introduction

Gastric cancer (GC) ranks third in cancer-related deaths worldwide^1^. The leading cause of death in GC is metastasis, in which most common form (50∼60%) is peritoneal metastasis (PM)^2,3^. Since GC patients with or without PM receive substantially different treatment strategies, including surgery and neoadjuvant therapies, accurate PM diagnosis is of great clinical significance for treatment and prognosis^4–6^.

The gold standard for PM diagnosis in GC is histopathological examination of peritoneal tissue biopsy, which is invasive and thus suggested to be obtained intraoperatively under laparoscopy. The procedure of histopathological diagnosis of PM is time-consuming and cannot provide results in a timely manner during surgery. Alternatively, preoperative computed tomography offers a way to detect PM noninvasively, but it is not sensitive enough, which leads to fatal false negatives^7,8^. Peritoneal lavage cytology, developed based on the theory that PM of GC is induced by colonization of exfoliated GC cells in the peritoneum, has been shown to be more sensitive than computed tomography, and more efficient and less invasive than histopathology^9^. Owing to these advantages, clinical guidelines in various countries have recommended peritoneal lavage cytology for patients with locally advanced GC during surgery^10–12^, which can be potentially extended to preoperative diagnosis and postoperative follow-up^13^. Nevertheless, the accuracy of conventional peritoneal lavage cytology for PM diagnosis is still limited, with sensitivity even lower than 60%^14–18^, and highly relies on pathologists^19^.

To increase the detection accuracy of GC cells in ascites, several biochemical and molecular biology methods have been developed, such as enzyme-linked immunosorbent assay (ELISA)^20^ or flow cytometry-based detection of specific proteins^21,22^, reverse transcriptase polymerase chain reaction (RT-PCR) or fluorescence in situ hybridization (FISH)-based detection of specific genes^23–28^. However, these methods are too time-consuming for intraoperative detection. More recently, a label-free optically induced electrokinetics microfluidic method was developed to efficiently separate GC cells from ascites of six patients with purity up to 71%^21^, but its performance on PM detection was not shown. Therefore, a new cytology method with both high accuracy and efficiency is urgently needed for detection of PM in GC.

Besides gene and protein expression, altered cell metabolism has been recognized as a hallmark of human cancers^29,30^. Cancer cells dysregulate metabolic pathways by high rates of lipid synthesis to support rapid growth^29,30^. GC cells have been found to accelerate lipid synthesis and reduce lipid hydrolysis^31,32^, which leads to increased accumulation of excessive lipids in lipid droplets (LDs)^31,32^. More importantly, dysregulated lipid metabolism has been shown to promote cancer metastasis^33^, including PM^34^. Particularly, as discovered by metabolomics, a variety of lipid molecules, such as triglycerides, sterols, fatty acids, could be used as biomarkers for PM of GC^35^.

For single cell molecular analysis, Raman spectroscopy is a commonly used label-free method. Several studies have demonstrated the potential of Raman spectroscopy in cytopathology for diagnosis of cancers, including cervical cancer, lung cancer, and oral cancer^36–38^. However, due to the weak spontaneous Raman signals, Raman spectroscopy-based cytology took up to 8 hours for a complete analysis without spatial information. With remarkably boosted Raman signals, simulated Raman scattering (SRS) microscopy is a desirable method of label-free and high speed molecular imaging at the single cell level^39,40^. In recent years, SRS microscopy with Raman tags^41^ has been widely used in the study of cancer metabolism^42^ and diagnosis^43^. For instance, Ji et al. for the first time employed two-color SRS microscopy to achieve virtual H&E staining, that is stimulated Raman histology (SRH)^44,45^, and later on Hollon et al. demonstrated deep learning-based SRH could realize intraoperative brain tumor diagnosis^46^. The SRH method has been applied in diagnosis of multiple human cancers^47–51^. Unfortunately, because cytology is not the gold standard for PM diagnosis in GC, the concept of SRH that primarily depends on morphology cannot be transferred to cytology-based PM diagnosis. Taken together, future cytology method of PM detection in GC probably requires an efficient acquisition of the information regarding both cellular morphology and composition, which is readily accessible by SRS microscopy.

In this study, we developed the stimulated Raman cytology (SRC), an SRS microscopy-based intelligent cytology method for diagnosis of PM from GC. By incorporating deep learning based single cell segmentation algorithms with three-color SRS imaging at the Raman bands corresponding to DNA, protein, and lipid, we first extracted 19 single cell features of morphology and composition from the exfoliated cells of ascites. Among them, 12 features including cellular area, lipid protein ratio, and lipid droplets number, etc. of exfoliated cells (N=53951 cells) were significantly different between PM positive and negative. Then, by newly developed hybrid K-means cell clustering and principal component analysis (PCA) algorithm (K-PCA), the differential raw features were dimensionally reduced and transformed to feature matrix in clustered latent space, which allowed us to find out the significant marker cell population. Finally, the feature divergence of significant marker cells was used to differentiate the PM positive and negative. A panel of machine learning classifiers, such as support vector machine (SVM), linear discriminate analysis (LDA), and logistic regression (LR), were used to train the PM diagnostic model with the feature matrix and the ground truth of PM results as inputs. By cross-validation, the sensitivity and specificity of our SRC method for PM detection were 81.5% and 84.9% respectively (n = 80 patients) within 20 minutes. Particularly, by providing composition information, the sensitivity of PM detection by the SRC significantly improved from 59.25% to 81.5%, suggesting that both cellular morphology and composition are essential for accurate diagnosis. Collectively, our SRC method may open up new opportunities for accurate and rapid detection of PM in GC with minimal invasion.

## Results

### Workflow of stimulated Raman cytology (SRC)

As shown in **Fig. 1**, the workflow of SRC is described. 1) Firstly, the three-color SRS imaging were performed on the exfoliated cells collected from the ascites of patients with locally advanced GC (**Fig. 1a**). Specifically, we acquired the SRS images of the Raman band for CH_2_ stretching in lipids around 2850 cm^-1^, the Raman band for CH_3_ stretching in proteins around 2930 cm^-1^ and the Raman band for CH_3_ antisymmetric stretching in DNA around 2965 cm^-1 52^, respectively. 2) Secondly, individual exfoliated cells were segmented based on the SRS images of DNA Raman band, by using deep learning based *Stardist* model (**Fig. 1b**). 3) Thirdly, a variety of features on cellular morphology and composition of single exfoliated cells for each patient were extracted. The features with statistically significant difference between PM positive and PM negative specimens were further selected to create the raw feature map (**Fig. 1c**). 4) Fourthly, the raw feature map then went through dimensionality reduction by principal component analysis (PCA) to get the latent space, including PC1, PC2 components etc. for each cell. The principal component was then transformed by K-means clustering to get transformed feature matrix in clustered latent space of PCA, such as PC1-center value, PC2-center value, etc. that were the medians of the corresponding PC values for each cell cluster (**Fig. 1d**). Here, we defined this hybrid PCA and K-means clustering algorithm as K-PCA, in which the PCA filtered out features with low standard deviation and K-means clustering built features spacing between clusters. The transformed feature matrix by K-PCA represented divergence of features for each cell cluster. The PC1 vs PC2 plot showed feature matrix of PC 1/2/3 values of Cluster #1/#2/#3 cells from PM positive and PM negative specimens (**Fig. 1d**). 5) Finally, the transformed feature matrix was used to train machine learning-based PM diagnostic models with histopathology as the ground truth. The performance of our SRC method in PM detection, such as sensitivity, specificity, and the area under receiver operating characteristic (ROC) curve, were further evaluated by leave-one-out cross-validation (**Fig. 1e**). The detailed procedures were described in the Materials and Methods section.

**Fig. 1.**
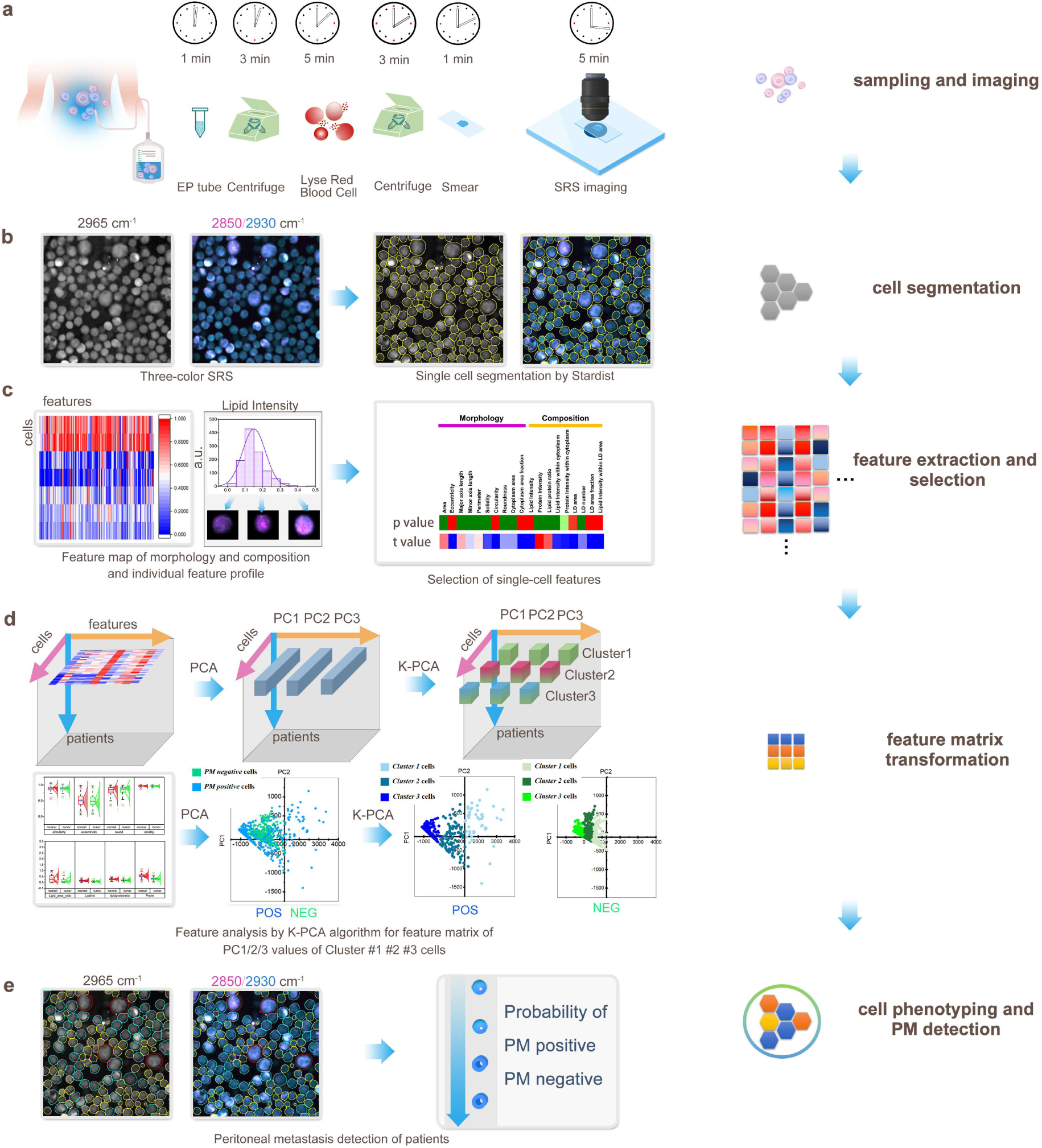
Workflow of stimulated Raman cytology (SRC). **a**, Sample preparation and SRS imaging. **b**, Single cell segmentation based on three-color SRS images. **c**, Single cell feature extraction and selection. **d**, Feature matrix transformation by K-PCA algorithm. **e**, Cell phenotyping by K-PCA and PM detection by machine learning classifiers.

### Single cell segmentation and feature extraction

By using deep learning *Stardist* network (**Fig. S1a**), we achieved single cell segmentation based on the SRS image of DNA Raman band around 2965 cm^-1^ with great performance (Dice parameter 0.89, IoU 0.81, and RRSE 2.3%), which was significantly better compared to conventional *watershed* or *flood fill* methods (**Supplementary Fig. S1b-c**). As shown in **Supplementary Fig. S2**, the feature extraction workflow was described in detail. The morphology features, including cellular area, minor axis length, major axis length, circularity, roundness, eccentricity, perimeter, and solidity, were extracted based on cell masks obtained from the 2965 cm^-1^ channel. The composition features, including lipid intensity, protein intensity, and lipid protein ratio, were extracted based on the intensity values obtained from 2850 cm^-1^ and 2930 cm^-1^ channels within the cell masks. According to the 2850 cm^-1^ channel, we recognized LDs by adaptive thresholding, which allowed quantification of LD area, LD number, LD area fraction, and lipid intensity within LD area. Based on calculations of 2850 cm^-1^ and 2965 cm^-1^ channels, nuclei could be identified by adaptive thresholding, which permit measurements of nucleus area, cytoplasm area, and cytoplasm area fraction. Then, the lipid intensity and protein intensity within cytoplasm could be quantified. The detailed procedures of feature extraction were described in the Materials and Methods section.

### SRC of gastric cell lines

Considering that peritoneal lavage exfoliated cells in GC patients are primarily composed of benign gastric epithelial cells, mesothelial cells, and malignant GC cells, we performed SRC on three presentative cell lines, including gastric epithelium GES-1, differentiated carcinoma SNU-16, and mesothelium HMrSV5. As shown in **Supplementary Fig. S3a-b**, we firstly extracted three features of cellular morphology (area, circularity, and roundness) and four features of chemical composition (lipid intensity, protein intensity, lipid protein ratio and LDs area fraction) for individual cells. Among these features, the lipid intensity was significantly greater in malignant cancer cells compared to normal epithelial and mesothelial cells, suggesting that malignant GC cells might accumulate more lipids than other types of exfoliated cells. In addition, the correlation coefficients were relatively high within morphology features (for instance, roundness and circularity) or within composition features (for instance, lipid intensity and protein intensity), but were relatively low between morphology and composition features (**Supplementary Fig. S3c**). By using LDA algorithm based on all the features, we could differentiate the GC cells from normal epithelium and mesothelium with the accuracy of 98.36% and 100%, respectively (**Supplementary Fig. S3d**). These results from cell lines demonstrated the potential of the SRC method for GC cell detection by integrating single-cell morphology and composition features.

### Differential features of cellular morphology and composition between PM positive and negative specimens

We quantitatively characterized 53951 individual exfoliated cells in ascites obtained from 80 GC patients (27 PM positive, 53 PM negative). As shown in **Supplementary Fig. S4**, 10 features of cellular morphology and 9 features of chemical composition were extracted. Among these features, 7 morphology features, including cellular area, major axis length, minor axis length, perimeter, solidity, cytoplasm area, and roundness, and 5 composition features, including protein intensity, lipid protein ratio, and lipid intensity within cytoplasm, protein intensity within cytoplasm, and LD number were significantly different between PM positive and PM negative specimens. As shown in the *T-value hot map*, the top three differential features were protein intensity, lipid protein ratio and cellular area. Nevertheless, the differences in composition and morphology features between PM positive and negative specimens were not very evident, which was likely due to the existence of heterogeneous populations in each specimen. Therefore, we proposed to further analyze the features by PCA and clustering algorithms.

### PM related significant marker cell population identified by K-PCA algorithm

Since exfoliated cells in PM positive specimens are composed of both tumor cells and normal cells (primarily mesothelial cells), it is necessary to identify the significant marker cell population with specific signatures for PM diagnosis. Firstly, the raw feature dataset was processed with dimensionality reduction by PCA, which produced the primary PC1 and PC2 components accounting for >90% of total features. Then, the feature components were transformed by K-means clustering to get transformed feature matrix of clustered latent space, including *Cluster1*-PC1, *Cluster1*-PC2, *Cluster2*-PC1, *Cluster2*-PC2, *Cluster3*-PC1, *Cluster3*-PC2, etc. *Cluster*-number defined the number of cells for each cluster. The principal components of cell features indicated three nearest neighbors by assessing the lowest neighbor distance. Therefore, we used three clusters for cell phenotyping. Without dimensionality reduction, the raw features were related to PM results with low correlation coefficient (R^2^ <0.3) (**Supplementary Fig. S5** and **Table S1**). With dimensionality reduction by K-PCA, the *Cluster1*-PC1 component was related to PM results with improved correlation coefficient (R^2^ = 0.67) (**Fig. 2a, Supplementary Fig. S5 and Table S2**). Based on the hot maps of raw features and feature matrix before and after K-PCA (n = 80 patients), *Cluster1* number and *Cluster1*-PC1 were obviously different between PM positive and negative (**Supplementary Fig. S5**). The number of cells in each cluster and the corresponding PC1 and PC2 values for totally 80 patients were shown in **Fig. 2a**. PC1-values of PM positive specimens were significantly higher than those of PM negative specimens (**Supplementary Table S2**). Particularly, the *Cluster1*-PC1 values of PM positive specimens were significantly greater than those of PM negative specimens, and a certain threshold could be used to differentiate PM positive and PM negative (**Fig. 2a**).

**Fig. 2.**
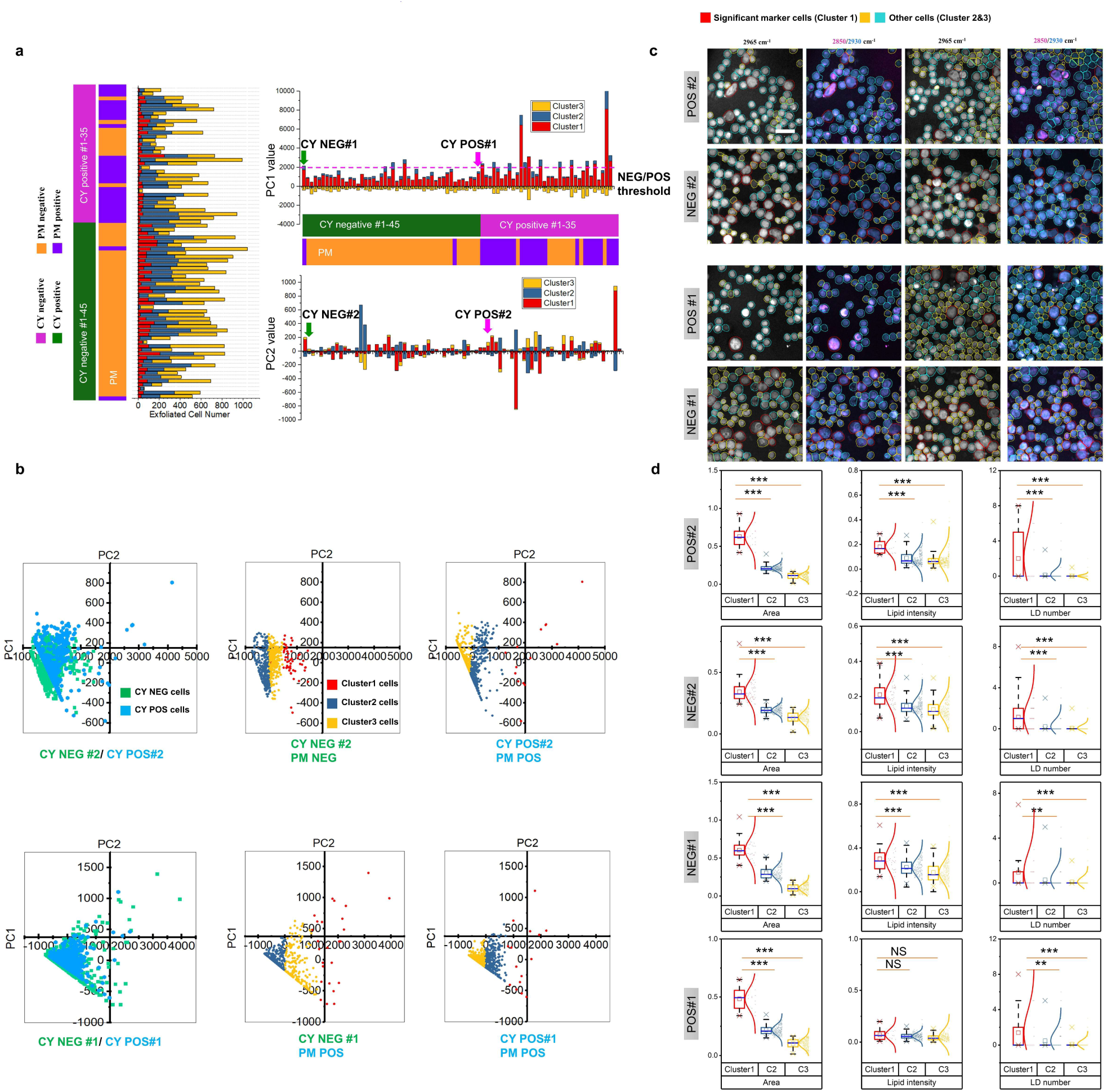
Feature matrix transformation by K-PCA algorithm. **a**, Exfoliated cell numbers, PC1 and PC2 values of clusters (*Cluster 1, 2, 3*) in latent space using PCA. **b**, Plot of PC1 vs PC2 for representative CY NEG#1 (diagnosed with PM positive), POS#1 (diagnosed with PM positive) and CY NEG#2 (diagnosed with PM negative), POS#2 (diagnosed with PM positive) before and after K-means clustering. **c**, Representative SRS images of CY NEG/POS#1 and CY NEG/POS#2. **d**, Quantitative comparisons of features (LD number, lipid intensity, cellular area) among clusters in CY NEG/POS#1 and CY NEG/POS#2. The box and whisker plots represent median values (center lines), mean values (horizontal bars), minimum and maximum (outliers), 25th to 75th percentiles (box edges) and 1.5x interquartile range (whiskers), with all points plotted. ***<0.0005, **<0.005, *<0.05, ns: no significant difference.

Notably, the *Cluster1*-PC1 values were more closely correlated with PM results than conventional cytology (CY) results of the same patients. For instance, the *Cluster1*-PC1 was higher than the threshold in CY negative#1 patient diagnosed with PM positive, but lower than the threshold in CY negtive#2 patient diagnosed with PM negative (**Fig. 2a**). By comparing the K-PCA latent spaces from two representative pairs of patients, as shown in **Fig. 2b**, the principal components of *Cluster1* from PM positive specimens dispersed to much greater distance within clusters relative to those from PM negative specimens, suggesting that *Cluster1* probably contained features closely related to PM. Thus, *Cluster1* was defined as the significant marker cell population as shown in **Fig. 2c**. The PC1 vs PC2 plot of CY negative#1 (diagnosed with PM positive) was indeed similar to those of PM positive. After cell clustering, we could characterize the morphology and composition features for different clusters of cells. As shown in **Fig. 2d** and **Supplementary Fig. S6**, morphology and composition features, such as cellular area, lipid intensity and LD number, were significantly different among different clusters. As shown in **Fig. 3**, we analyzed differences of features between all the PM positive and negative specimens. The overall 10 features such as cellular area, perimeters, cytoplasm area, cytoplasm area fraction, lipid intensity, protein intensity, lipid protein ratio, protein intensity within cytoplasm, lipid intensity within cytoplasm and LD number were significantly different between PM positive and negative after K-PCA (**Fig. 3a**). In terms of different clusters, the features, such as area, perimeter, cytoplasm area fraction, cytoplasm area and LD number, were significantly different between significant marker cells (*Cluster 1*) and other cells (*Cluster 2&3*) (**Fig. 3a**). These results together indicate that our K-PCA algorithm has a potential to accurately evaluate PM by introducing more information on composition besides morphology used in conventional cytology.

**Fig. 3.**
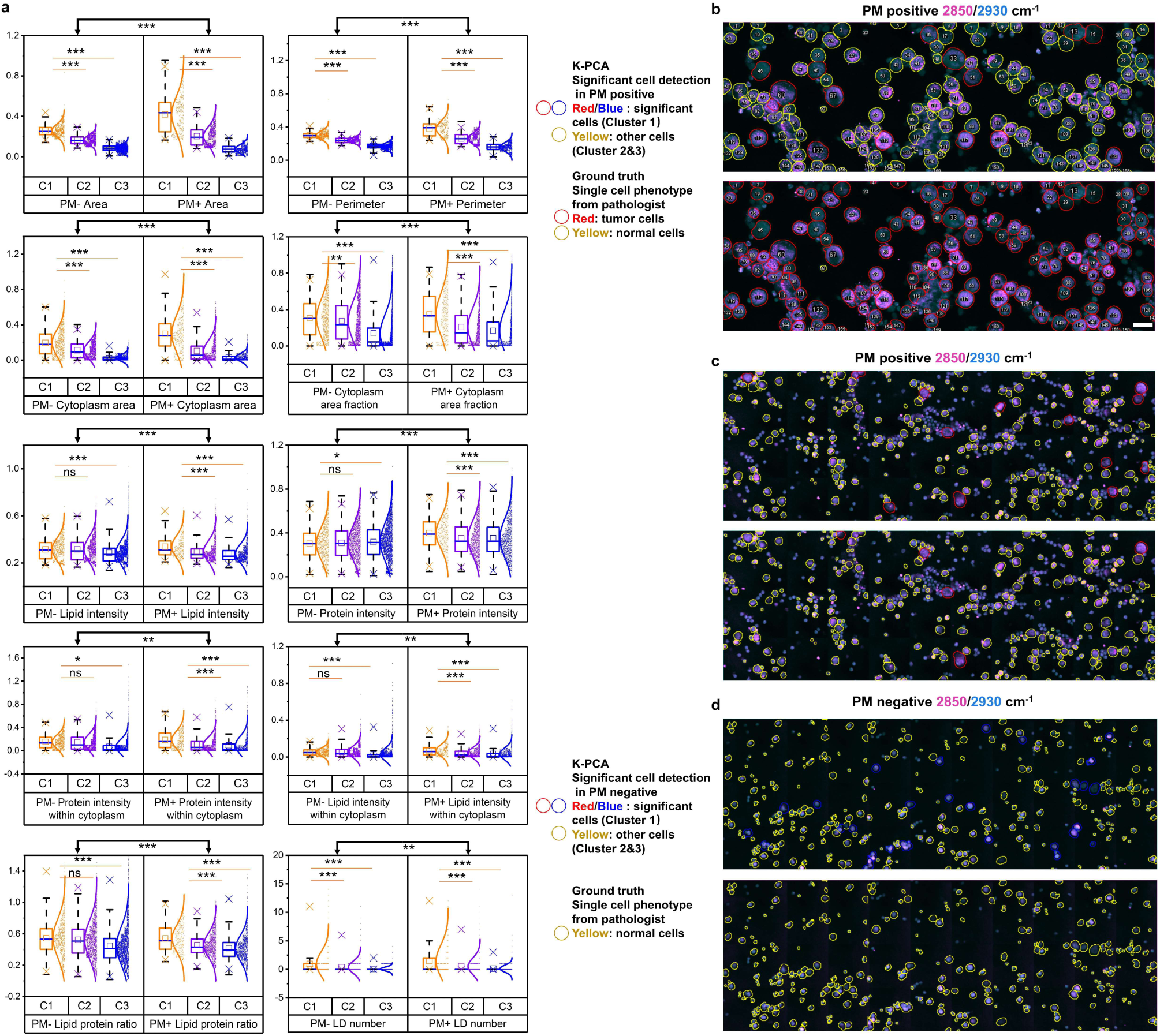
Significant marker cell identification by K-PCA algorithm. **a**, Quantitative comparisons of features among clusters (*Cluster 1*: significant marker cells; *Cluster 2&3*: other cells) between all PM positive and negative specimens. The box and whisker plots represent median values (center lines), mean values (horizontal bars), minimum and maximum (outliers), 25th to 75th percentiles (box edges) and 1.5x interquartile range (whiskers), with all points plotted. ***<0.0005, **<0.005, *<0.05, ns: no significant difference. **b-d,** Representative SRS images with clusters identified by K-PCA algorithm and tumor/normal cells identified by pathologists using the conventional cytology of the same specimen as reference, for PM positive specimen with high percentage of tumor cells (**b**), PM positive specimen with low percentage of tumor cells (**c**), and typical PM negative specimen without tumor cells (**d**). Scale bar, 20 µm.

Moreover, we explored the relationship between significant marker cells and tumor or normal cells. The SRS images of exfoliated cells were firstly stitched into larger scale (**Supplementary Fig. S7**) that could be comparable with the H&E cytological images of the same locations *in situ*. Based on the labeling of normal and tumor cells on the cytological images by pathologists, we then integrated machine learning (ML) methods with PCA, called ML-PCA here, to differentiate tumor cells from normal cells. The accuracy, sensitivity, specificity and the area under ROC curve (AUC) of ML-PCA methods were 93.8%, 94.1%, 93.6%, and 0.98 for SVM based ML-PCA, and 90.5%, 92.0%, 90.1%, and 0.96 for LDA based ML-PCA (**Supplementary Fig. S8**). As shown in **Fig. 3b-d** and **Supplementary Fig. S9**, the significant marker cell population in PM positive was primarily included in the tumor cell population, especially for PM positive specimens with high percentage of tumor cells. In the meanwhile, the significant marker cell population in PM negative specimen belonged to the normal cell population. These findings suggest that the significant marker cell population possibly contains features in which the divergence highly correlates with PM. The differences of features between significant marker cells and other cells identified by K-PCA were more evident than the differences of features between tumor cells and normal cells identified by ML-PCA, especially for PM positive specimens with high percentage of tumor cells (**Supplementary Fig. S10**). Collectively, in SRC method, we do not need to follow conventional cytology to capture all tumor cells, but rather use K-PCA algorithm to identify the significant marker cells with specific morphology and composition features, in which the divergence is likely related to PM.

### Demonstration of PM detection by SRC

The transformed feature matrix was further used as input features in machine learning models for PM prediction. As shown in **Fig. 4a**, the feature matrix obtained by K-PCA (*Cluster1*-PC1, *Cluster1*-PC2, *Cluster1*-number etc.) could differentiate PM positive and negative more clearly than the feature matrix obtained by ML-PCA. The feature matrix was then trained and validated with the ground truths of PM diagnosis by using machine learning methods, including SVM, LDA, and LG. With leave-one-out cross-validation, PM positive probability of each patient, ROC curve, and confusion matrix were shown in **Fig. 4b** and **Supplementary Fig. S11** (details in **Supplementary Table S3**). In terms of supervised ML-PCA method, SVM performed better than LDA and LG for differentiation between PM positive and PM negative, with AUC of 0.797, 63.0% sensitivity and 88.7% specificity at an appropriate cutoff (**Fig. 4b**). The results of LDA and LG classifiers for ML-PCA were shown in **Supplementary Fig. S11**. Compared to supervised ML-PCA, un-supervised K-PCA gained much better performance for PM detection, with AUC of 0.85, 81.5% sensitivity and 84.9% specificity at an appropriate cutoff, by using LG classifier (**Fig. 4b**). The results of SVM and LDA classifiers for K-PCA were shown in **Supplementary Fig. S11**. As shown in **Fig. 4c**, the PM prediction results based on ML-PCA and K-PCA for each patient were delineated. The directly visualized comparisons with the ground truths further demonstrated the great performance of K-PCA based PM detection. These results suggest that our SRC method is capable to detect PM with high sensitivity and specificity.

**Fig. 4.**
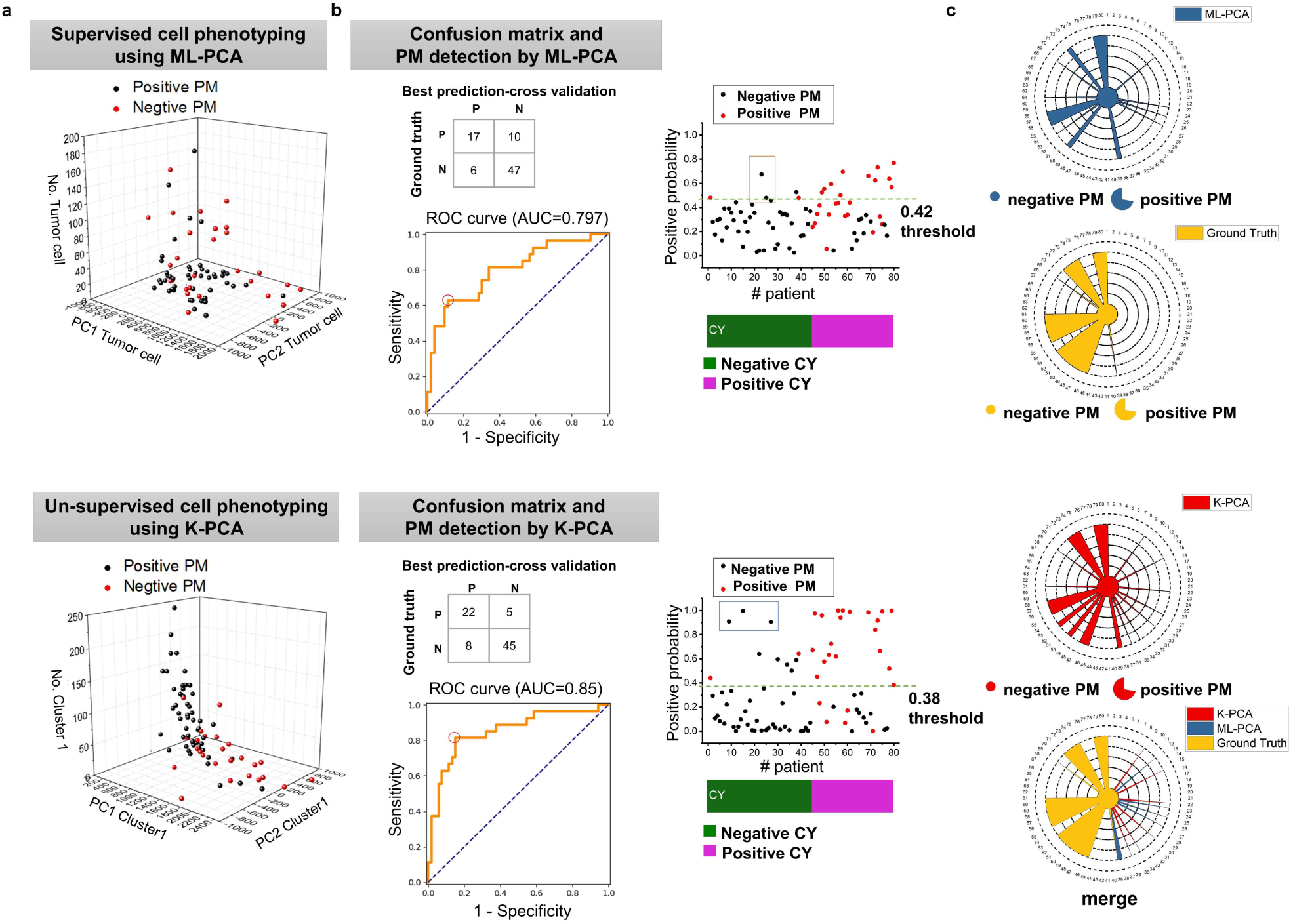
PM detection by SRC using machine learning classifiers. **a**, Upper: Input feature matrix (PC1, PC2, cell number of tumor cell) of PM positive and negative specimens by the ML-PCA method (SVM). Down: Input feature matrix (PC1, PC2, cell number of *Cluster 1*) of PM positive and negative specimens by the K-PCA method. **b**, Upper: Confusion matrix, positive probability with cross-validation, and ROC curve by the ML-PCA method (SVM). Down: Confusion matrix, positive probability with cross-validation, and ROC curve by the K-PCA method. **c**, Upper: Final detection results by ML-PCA (SVM) compared with ground truth. Down: Final detection results by K-PCA compared with ground truth.

### Interpretation of feature contribution in PM detection by SRC

In order to interpret the result by SRC, we analyzed importance coefficient of the single-cell features (**Fig. 5**). The diagnostic performance of SRC using both morphology and composition features was significantly better than that using either morphology or composition features. Specifically, the sensitivity increased from 59.25% to 81.5%, the accuracy increased from 76.25% to 83.75%, the specificity increased from 75.48% to 84.9%, negative predictive value (NPV) increased from 80.35% to 90%, and positive predictive value (PPV) increased from 62.85% to 73.3% (**Fig. 5a**). The improved sensitivity was predominantly contributed by composition features, whereas the improved specificity was mainly contributed by morphology features. In terms of NPV and PPV, both morphology and composition features contributed.

**Fig. 5.**
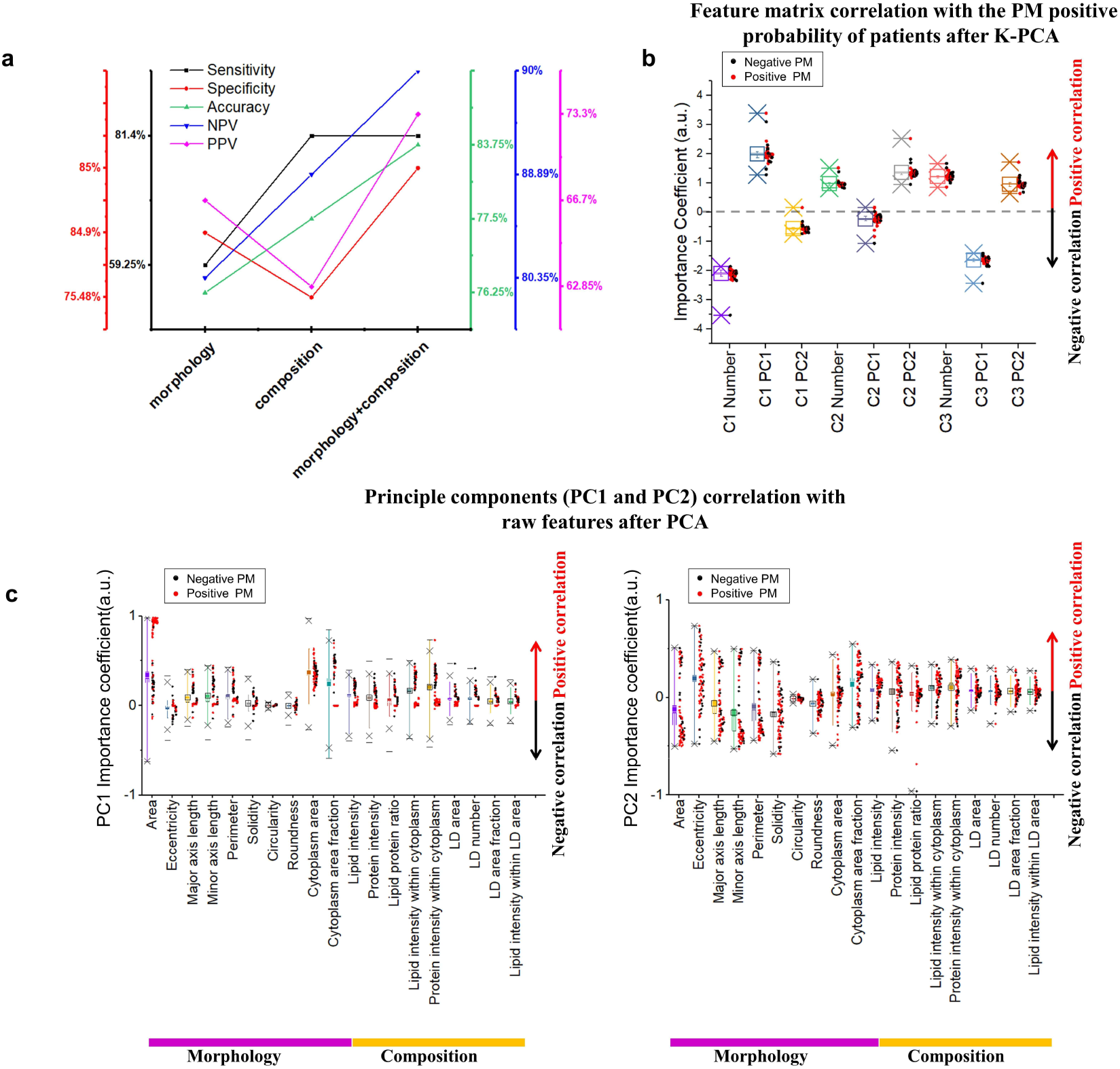
Feature contribution in PM detection by SRC. **a**, Comparisons of the performance of PM detection (sensitivity, specificity, accuracy, NPV and PPV) based on either or both of morphology and composition features. **b**, Importance coefficients of feature matrix to PM detection results. Feature matrix: C1 PC1: PC1 values of *Cluster1*, C1 Number: *Cluster#1 cell number*, etc. **c**, Importance coefficients of raw features to principal components, PC1 value (left) and PC2 value (right). The box and whisker plots represent median values (center lines), mean values (horizontal bars), minimum and maximum (outliers), 25th to 75th percentiles (box edges) and 1.5x interquartile range (whiskers), with all points plotted.

Moreover, we analyzed the importance coefficients of feature matrix, which contributed to final PM results (**Fig. 5b**). Importance coefficients represent the relationship between the given feature matrix and PM results, assuming that all the other matrix components remain constant (conditional dependence)^53^. Through analyzing importance coefficient of feature matrix by K-PCA, the *Cluster1*-number was negatively related with PM results and *Cluster1*-PC1 was positively related with PM results, suggesting that clustering benefit PM detection (**Fig 5b**). We also quantitively analyzed the importance coefficients of raw features, suggesting the correlation between principal components and raw features. As shown in **Fig. 5c**, principal components (PC1 and PC2) had the top three highest correlations with cellular area, cytoplasm area fraction, and protein intensity within cytoplasm. The features, including cell area, cytoplasm area, lipid intensity and protein intensity within cytoplasm etc., had positive correlations with PC1 (**Fig. 5c**); whereas the features, including cytoplasm area, cytoplasm area fraction, lipid intensity and protein intensity within cytoplasm etc., had both positive and negative correlations with PC2 (**Fig. 5c**). Therefore, our quantitative interpretation demonstrates that morphology and composition features of single exfoliated cells after clustering contribute to accurate PM detection by the SRC method.

## Discussion

In this study, we have developed the SRC, a label-free SRS microscopy-based intelligent cytology method, for detection of PM in GC. Integration of three-color SRS microscopy with deep learning segmentation model provided both morphology and composition features of single cells. Our hybrid PCA and K-means clustering analysis further transformed these raw features to feature matrix and enabled identification of the significant marker cell population, the divergence of which was strongly related to PM. The transformed feature matrix was finally used as input features in machine learning models for PM prediction. For 80 GC patients, the SRC achieved 81.5% sensitivity, 84.9% specificity, 83.75% accuracy, and AUC of 0.85, by using cross-validation compared with histopathology, the gold standard of PM detection. Moreover, the average test time for each patient was within 20 minutes. As discussed below, our study demonstrates that the SRC may open up a new avenue for accurate and rapid detection of PM from GC.

Firstly, our SRC method enhances accuracy of PM detection for GC by integrating morphology features and composition features of single exfoliated cells. Conventional peritoneal lavage cytology entirely relies on cell morphology, which reaches specificity as high as 85% but limits the sensitivity to even lower than 60% for PM diagnosis^16^. This suggests that the morphology feature largely contributes to the specificity but is not enough to gain the sensitivity. Thus, the concept of well-established SRS-based virtual histology (SRH) method that primarily depends on morphology could not be transferred to cytology. Different from conventional cytology and SRH, our SRC brought in composition features, which indeed significantly increased the sensitivity from 59.25% to 81.5%. Particularly, lipid protein ratio, lipid intensity within cytoplasm, and protein intensity within cytoplasm were recognized to be the most critical composition features contributing to sensitive PM detection.

Secondly, the performance of our SRC method depends on accurate single cell phenotyping, which combines single cell segmentation and cell phenotyping. The single cell segmentation was conducted by using pre-trained deep learning *Stardist* model to achieve high accuracy, especially for those closely touching cells that were difficult to be separated by common segmentation algorithms. The single cell phenotypes were further recognized by newly developed hybrid K-PCA algorithm based on feature dimensionality reduction and clustering. Importantly, with specific morphology and composition features, we identified the significant marker cell cluster, the divergence of which was then used to differentiate the PM positive and negative. Similarly, a recent work demonstrated the existence of significant marker cell population at the edge between tumor and stroma for accurate prediction of tumor response to immunotherapy^54^. The above evidence collectively indicates that the significant marker cells with specific molecular signatures support accurate detection of PM from GC.

Thirdly, our SRC method using unsupervised K-PCA is significantly more precise than supervised ML-PCA for PM detection. We made “cell to cell” inspections on SRC and conventional cytology of the same specimen and classified normal/tumor cell with 93% accuracy. The 7% error of ML-PCA based cell classification may induce bigger error for PM detection. The feature matrix transformed by K-PCA was used to diagnosis PM with much higher accuracy. With the leave-one-out cross-validation of 80 patients, LG showed the best performance for PM diagnosis among three diagnostic models (SVM, LDA and LG) (**Supplementary Table S4**). The detailed explanation was described in **Supplementary Note 1**.

Fourthly, we further investigated the mismatched results of SRC compared with the gold standard histopathology. As shown in **Supplementary Table S5**, the results of SRC were close to histopathology but not conventional cytology. Compared to histopathology, SRC had 5 false negatives out of 80 patients. In order to find out the possible reason of false negative, we compared SRC with conventional cytology on the same specimen *in situ*. As shown in **Supplementary Fig. S12**, different from the original positive cytology result, the cytology result obtained from the same specimen analyzed by SRC showed negative result (i.e. no tumor cells). Similarly, as shown in **Supplementary Fig. S13**, different from the original negative cytology result, the cytology result obtained from the same specimen analyzed by SRC showed positive result. These results suggest that the false negative in SRC is possibly related with the throughput of our method and may occur in the specimen with very low percentage of tumor cells. In addition, compared to histopathology, SRC had 8 false positives out of 80 patients, which may be due to limited features obtained from three-color SRS imaging. The sensitivity and specificity of SRC could be further improved by increasing throughput, for example via stimulated Raman flow cytometry^55,56^, and by introducing more composition features, for example via hyperspectral SRS imaging cytometry^57^.

Finally, our SRC method may enable a variety of clinical practices for better management of locally advanced GC. For instance, besides rapid detection of PM during surgery, SRC could also facilitate preoperative diagnosis and postoperative follow-up, which can be hardly done by histopathology due to its invasiveness. As shown in **Supplementary Fig. S14**, we tried to assess the changes of PM upon chemotherapy after surgery and predict prognosis. Moreover, besides PM detection, cytology itself is also a very important indicator for potential PM that may have not occurred yet. Fortunately, the SRC is based on multi-color SRS microscopy that has been previously established to permit virtual H&E staining^46,47^. Thus, the integration of SRC with virtual staining may be desirable to give diagnostic results of both PM and cytology. Furthermore, the concept of SRC could be extended to other clinical scenarios of cancer diagnosis using cytology, such as Pap smear for cervical cancer, urinary cytology for urothelial cancer, bronchoalveolar lavage fluid examination for lung cancer, etc. Taken together, SRC may hold great promise for becoming the next generation of cytology in the near future.

## Methods

### Cell culture

GES-1, HMrSV5, SNU-16 cells were cultured in RPMI 1640 (Gibco, 11875119), supplemented with 10% fetal bovine serum (Omega Scientific, FB-21), and 0.1% penicillin/streptomycin (Gibco, 15070063). Cell cultures were incubated in an incubator at 37 °C with 5% CO_2_.

### Clinical sample

Peritoneal lavage fluids were collected from 80 patients diagnosed with locally advanced GC in the Peking University Cancer Hospital, with 27 PM positive, 53 PM negative; and 35 CY positive, 45 CY negative. The detailed sex, age, PM positive/negative, and CY positive/negative of each patient were described in the **Supplementary Table S3**. For each patient, half (100mL) of the sample was inspected by conventional cytology by pathologists, and the other half (100mL) was analyzed by the SRC method. This study was approved by the Institutional Review Boards of Peking University Cancer Hospital and Beihang University.

### Diagnostic laparoscopy staging and conventional cytology examination

Intraoperative laparoscopy was performed under general anesthesia. The patient was placed in a supine position. A 10-mm disposable trocar (observing hole) was inserted into the sub-umbilicus, and a 30° telescope was used. Another 10-mm trocar and a 5-mm trocar were inserted into the right and left upper quadrants, respectively. Prior to any manipulation, 250 mL of warm normal saline was infused into the subphrenic space, subhepatic space, omentum, bilateral paracolic sulci and the pouch of Douglas. Care was taken to avoid direct contact of the irrigation with the primary tumor. At least 200 mL of fluid was aspirated from the subphrenic space, subhepatic space and pouch of Douglas. The fluid was immediately sent for the SRC and cytological examination. For conventional cytology, the exfoliated cells were stained with hematoxylin and eosin (H&E). Two professional pathologists examined the H&E slides independently. Patients with negative cytology results were labeled as CY–, and patients with positive cytology results were labeled as CY+. Subsequently, a systematic inspection of the abdominal cavity was performed clockwise from the right quadrant. Any suspicious lesion was biopsied and sent for histopathologic examination. Patients with negative histopathologic results were diagnosed as PM negative (PM–), and patients with positive histopathologic results were diagnosed as PM positive (PM+).

### Preparation of the exfoliated cell for SRS imaging

The ascites samples were first centrifuged at 2000 rpm for 3 minutes. Then, the concentrated samples were treated by red blood cell lysate for 5 minutes and rinsed with phosphate buffered saline. The sample (∼10 μL) was dropped on the glass slide and evenly smeared in an area of about 1 cm^2^. The slide was used for SRS imaging after air-drying and stored at -80°C for future use if needed.

### Three-color SRS microscopy

Our SRS microscope employed a dual output picosecond pulse laser (picoEmeraldTM S, Applied Physics & Electronics) with a repetition rate at 80 MHz and 2 picosecond width. The laser has an integrated output for both the pump beam with tunable wavelength from 700 nm to 960 nm and the Stokes beam with fixed wavelength at 1031 nm, which are overlapped in space and time. When performing SRS, the Stokes beam was modulated at ∼20 MHz by an electronic optic modulator. The collinear pump and Stokes beams were coupled to a two-dimensional scanning galvanometer (GVS012-2D, Thorlabs) and then imported into an inverted microscope (IX73, Olympus). A 60X water immersion objective lens (LUMPlanFL N, 1.0 numerical aperture, Olympus) focused the lasers into the sample. The photons were collected by an 60X water condenser (LUMPlanFL N, 1.0 numerical aperture, Olympus). The pump beam was selected by a short-pass filter (ET980SP, Chroma), and was detected by a photodiode (S3994-01, Hamamatsu, Japan) equipped with a resonant circuit that selectively amplifies the signal at the optical modulation frequency. The stimulated Raman loss signal was then extracted by a digital lock-in amplifier (HF2LI, Zurich Instrument, Zurich, Switzerland). The output voltage from the lock-in amplifier, which represents the SRS signal, was sampled by a DAQ card (PCI-6363, National Instruments, Austin, TX). A LabVIEW platform synchronized scanning of wavelength with 2D multivariable acquisition of XY images. SRS imaging was performed by tuning 796.8nm, 791.8nm and 789.5nm to get the Raman band of 2850 cm^-1^, 2930 cm^-1^, and 2965 cm^-1^, which correspond to the C-H stretching region of lipid, protein, and DNA, respectively. The excitation power at the sample was ∼20 mW for pump and ∼90 mW for Stokes for all cell samples. Each SRS image contained 400×400 pixels. The pixel dwell time was 10 μs. The field of view of single SRS images was 100×100 μm.

### Image processing and analysis

For SRS image processing, home-built Python programs were used to segment single cell with masks, extract features and classify single cell with labelled masks. The whole process was described in the supplementary **Supplementary Table S6**.

The single cell segmentation ran with the Python library *stardist2D 0.7.1* under *tensorflow 1.7*. The feature extraction ran with the Python library *skimage 0.18.0*. The cell phenotyping and dimensional reduction using K-PCA and ML-PCA ran with the Python library *sklearn 0.21.2*. PCA as a linear dimensionality reduction uses singular value decomposition (SVD) of the features data to project it to a lower dimensional space. The input feature data is centered but not scaled for each feature before applying the SVD. The K-means algorithm clusters input features data by separating cells in *n* groups of equal variances, through minimizing a criterion known as sum of squares within cluster.

After cell phenotyping, labelled masks were created and saved with virtual pseudo colors. The saved ROI files could be opened by the *ImageJ*. The significant feature components (cellular area, perimeter, cytoplasm area, lipid intensity, protein intensity, and lipid protein ratio etc.) of three clusters were saved in csv files. Imported with the significant components, the supervised classifiers (SVM, LG and LDA) all ran with the Python library *sklearn 0.21.2*.

The positive probability of PM, confusion matrix and ROC curve for each of the 80 patients were plotted and saved. The whole image processing and analysis (**Supplementary Table S6**) for each patient with 500 cells took less than 1 min on a customized computer equipped with a single graphics card (EVGA GeForce GTX 2080Ti FTW3) and an Intel i9-7920x CPU.

### Large-scale SRS imaging

We programmed a home-built code for large-scale SRS imaging including automatic scanning, automatic stitching, and automatic calibration. Automatic scanning was programmed by home-built LabVIEW software. We acquired 4*10 images with 20% overlay to create a FOV of 400*1000 μm^2^. After image acquisition, the stitching algorithm programmed by MATLAB automatedly calibrated and smoothed the edges between images. The shifts among three-color SRS image channels were corrected. The schematic and typical results were described in the **Supplementary Fig. S7**.

### Extraction of features from single cells

We totally extracted 19 features, including morphology and composition, of single exfoliated cells, as shown in **Supplementary Fig. S2**. The morphology features included area, eccentricity, major axis length, minor axis length, perimeter, solidity, circularity, roundness, cytoplasm area fraction, and cytoplasm area. Major and minor axis length are the primary and secondary axis of the best fitting ellipse with the selection cell contour. The solidity is convex area of the convex hull that encloses single cell divided by cellular area. The eccentricity is minor axis length divided by major axis length. The circularity and roundness are calculated by the following equations, respectively.

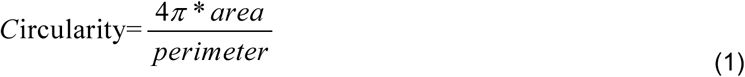

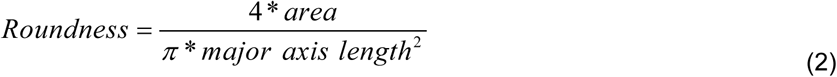

The composition features included lipid intensity, protein intensity, lipid protein ratio, lipid intensity within cytoplasm, the protein intensity within cytoplasm, lipid droplet (LD) area, LD number, LD area fraction, and lipid intensity within LD area. The lipid and protein intensity of each image were calculated from the intensity of SRS image at 2850 cm^-1^ and 2930 cm^-1^ by the following formula.

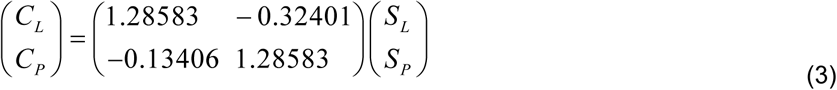

S_L_ is the signal intensity of SRS image at 2850 cm^-1^, S_P_ is the signal intensity of SRS image at 2930 cm^-1^, C_L_ is the lipid intensity and C_P_ is the protein intensity of each image. The constants were measured from pure BSA and oil by SRS imaging at 2930 cm^-1^ and 2850 cm^-1^.

As shown in **Supplementary Fig. S2**, the 8 morphology features such as area, eccentricity, major axis length, minor axis length, perimeter, solidity, circularity and roundness were extracted from cellular masks by using *skimage.measure* function in Python script as same as the ‘Measure’ module at ImageJ after single cell segmentation.

Then, the lipid intensity, protein intensity, and lipid protein ratio of single cells were also extracted using *skimage.measure* function based on C_L_, C_P_ and cellular masks. Then, nucleus segmentation masks were extracted from DNA channel by adaptive thresholding. By using *skimage.measure* function, cytoplasm area were calculated by cytoplasm masks, which are cell masks subtracted by nucleus masks. Cytoplasm area fraction is cytoplasm area divided by cell area. Then another 2 composition features (lipid intensity within cytoplasm, and protein intensity within cytoplasm) were calculated by C_L_, C_P_ and cytoplasm masks.

Finally, we got masks of LD area from 2850 cm^-1^ by Python library *cv2.threshold*, as same as the ‘Threshold’ module at ImageJ. LD in single cells can be segmented due to their higher local signal intensities compared to surrounding cellular compartments. By using *numpy.percentile* function in Python script, we could get intensity threshold value of each image from the overall intensity histogram. Then based on LD masks, we extracted the LD number and LD area. LD area fraction is LD area divided by cell area. Lipid intensity within LD area is calculated by C_L_ and LD mask. The segmented LDs areas were highly consistent with visual judgment as shown in **Supplementary Fig. S2**.

### Quantitative evaluation of single cell segmentation

Dice parameter, relative mean square error (RMSE) between ground truth and automate cell segmentation were used to quantitatively assess the performance of cell segmentation. Dice parameter represents the overlap of segmentation area by *stardist* 2D and visual judgment by the formula.

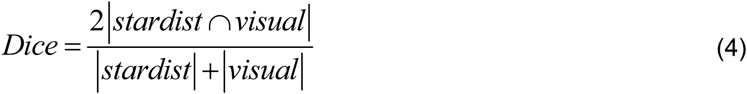

RMSE represents relative mean square error of cell counting between *stardist* 2D and visual judgment in each image.

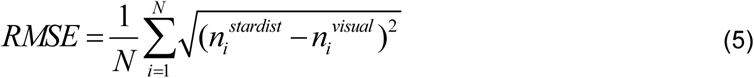

n_i_^stardist^ means cell counting number by *stardist* model and n_i_^viusal^ means cell number from visual judgement in each image *i*.

### Statistical analysis

The box and whisker plots with Origin 2017 represent median values (center lines), mean values (horizontal bars), minimum and maximum (outliers), 25th to 75th percentiles (box edges) and 1.5x interquartile range (whiskers), with all points plotted. Student’s t test was used for comparisons between groups, and p < 0.05 was considered statistically significant. T value and p value were performed by Origin 2017 using Student’s t test. The positive and negative classification for PM was evaluated with ROC curve analysis. The area under the ROC curve (AUC) ranging from 0 to 1 evaluates the ability of a model to accurately distinguish two categories.

### Reporting Summary

Further information on research design is available in the Nature Research Reporting Summary linked to this article.

## Data availability

The main data supporting the findings of this study are available within the paper and its Supplementary Information. The training dataset and the output images for Stardist2D network for single cell segmentation is available at https://doi.org/10.48550/arXiv.1806.03535. All data for research purposes are available in the main text or the supplementary materials upon reasonable request.

## Code availability

The original code for Stardist2D is available at https://github.com/qupath/models/tree/main/stardist. We apply this code to our dataset. MATLAB was used for creating image tiles for the network and for restitching the output image tiles. All the programs for research purposes are available upon reasonable request. The system control software and the data collection software are proprietary and used in licensed technologies.

## Author contributions

Conceptualization: S.H.Y., Z.Y.L., Z.Q.W., Y.B.F., J.F.J., Methodology: X.C., S.H.Y., Z.Q.W., Specimen: Q.W., Z.Q.W., Z.W.L., F.S., Experiment: X.C., Z.H., Y.X.H., W.H.Z., K.J.Z., Q.W., Algorithm: X.C., Z.Q.W., K.J.Z., Funding acquisition: S.H.Y., X.C., Z.Q.W., Y.B.F., Project administration: S.H.Y., Z.Q.W., X.C., P.W., Supervision: S.H.Y., Z.Y.L., Y.B.F., J.F.J., Writing – original draft: X.C., Z.Q.W., S.H.Y., Writing – review & editing: S.H.Y., X.C., Z.Q.W., Y.B.F., Z.Y.L., J.F.J..

