## Supporting information for "Accurate and Rapid Detection of Peritoneal Metastasis from Gastric Cancer by AI-assisted Stimulated Raman Cytology"

[1]Key Laboratory of Biomechanics and Mechanobiology (Beihang University), Ministry of Education, Institute of Medical Photonics, Beijing Advanced Innovation Center for Biomedical Engineering, School of Biological Science and Medical Engineering, Beihang University, Beijing 100083, China
[2]Gastrointestinal Cancer Center, Key Laboratory of Carcinogenesis and Translational Research (Ministry of Education), Peking University Cancer Hospital & Institute, Beijing 100142, China
[3]School of Engineering Medicine, Beihang University, Beijing 100083, China
[4] Key Laboratory of Carcinogenesis and Translational Research (Ministry of Education), Department of Pathology, Peking University Cancer Hospital & Institute, Beijing 100142, China
[&]These authors contributed equally.

**Supplementary Materials**

Supplementary Note 1: Cell phenotyping with SRC and H&E cytology of the same sample

Supplementary Fig. S1. Performance of single cell segmentation by stardist model.

Supplementary Fig. S2. Workflow of feature extraction based on masks.

Supplementary Fig. S3. Results of SRC of gastric cell lines with single cell segmentation and classification.

Supplementary Fig. S4. Representative SRS imaging of GC exfoliated cells and comparisons of 19 features between positive and negative PM.

Supplementary Fig. S5. Hot map of features raw feature and feature matrix transformed by K-PCA.

Supplementary Fig. S6. Representative images and quantitative analysis of exfoliated cell phenotyping.

Supplementary Fig. S7. Image stitching of three-color SRS imaging of exfoliated cells.

Supplementary Fig. S8. In situ SRS&HE imaging of exfoliated cells.

Supplementary Fig. S9. Typical tumor cell and normal cell detection by ML-PCA.

Supplementary Fig. S10. Results of feature profiles by K-PCA and ML-PCA.

Supplementary Fig. S11. Results of PM detection by classifiers (SVM, LDA, LG etc.).

Supplementary Fig. S12. Typical SRS&HE in-situ images for SRC negative detection.

Supplementary Fig. S13. Typical SRS&HE in-situ images for SRC positive detection.

Supplementary Fig. S14. PM positive probability before and after chemotherapy for patients diagnosed with PM positive (n = 3) and PM negative (n=2).

Supplementary Table S1. Typical raw features of 80 patients.

Supplementary Table S2. Values of Cluster1-PC1, Cluster1-PC2 and Cluster2-PC1, Cluster2-PC2 of 80 patients.

Supplementary Table S3. Diagnostic results for each patient, including conventional cytology, histopathology, and SRC (based on SVM, LDA, or LG models) predicted probability of PM.

Supplementary Table S4. Performance comparisons between K-PCA and ML-PCA methods.

Supplementary Table S5. Comparisons between our SRC results and conventional cytology, histopathology, respectively.

Supplementary Table S6. Image acquisition and processing workflow by task, software, and description.

**Supplementary Note1: Cell phenotyping with SRC and H&E cytology of the same sample**

We compared the performance of cell phenotypes algorithms using in situ SRS&HE cytology. We firstly imaged the exfoliated cells using Stimulated Raman cytology. Then we labeled the samples with H&E dye and imaged in situ cells of same slices. Using SRC method, we stitched 4*10 FOVs to create a large scope comparable with traditional H&E cytology with a customized MATLAB program. The imaging results of the label-free exfoliated cells were compared with cytological images in situ with hematoxylin and eosin (H&E) staining by “cell to cell” (**Supplementary Fig. S8**). Based on the single cell labelling from pathologists, we built a machine learning based model (LDA and SVM) to classify normal/tumor cells. The training and test dataset (8:2) include total 12620 cells (1158 normal cells; 11462 tumor cells). The best accuracy of single cell classification was could be 93.8% after our model optimization.

We also studied the cell phenotype algorithms for PM detection using K-PCA and ML-PCA (**Supplementary Fig. S9 and Fig. 3**). Using K-PCA algorithm, we could filter out significant marker cells from bundle of normal/tumor cells (**Supplementary Fig. S9 and Fig. 3**). For PM positive with lots of tumor cells, the composition features (LD number) and shape features (cellular area and cytoplasm area fraction) of significant marker cells (*Cluster 1*) were significantly different than other tumor cells (*Cluster 2&3*) (**Supplementary Fig S10**). For PM positive with few tumor cells, the feature divergence between clusters becomes weaker. Meanwhile, cellular area, lipid intensity, cytoplasm area fraction and LD number after ML-PCA become more significantly different than K-PCA. Additionally, the lipid intensity, LD number of tumor cells are higher than normal cells, which are consistent with previous studies[1]. For PM negative without tumor cells, cellular area, LD number, and cytoplasm area fraction of significant maker cells (*Cluster 1*) were significantly different than other normal cells (*Cluster 2&3*) (**Supplementary Fig S10**).

We also compared the features of shape, composition especially LDs, and the raw features were mixed between positive/negative PM before K-PCA. After K-PCA based dimensional reduction, the positive/negative PM were obviously separated with *Cluster1*-number and *Clutser1*-PC1 (**Supplementary Fig. S5**). The final PM detection results from cell phenotype were shown in **Fig. 4** and **Supplementary Fig. S11**. Incorporating with LDA, SVM and LG classifiers, the unsupervised K-PCA for PM detection was better than supervised ML-PCA with leave-one-out cross-validation. Specially, the sensitivity of unsupervised K-PCA method performs better (**Supplementary Fig. S11** and **table S4**). The final PM detection accuracy of K-PCA with LG, SVM and LDA were respectively 83.75%, 78.75% and 77.5%%; ML-PCA were 80%, 72.5% and 76.25% respectively as shown in **Fig. 4** and **Supplementary Fig. S11**.

By using SRS&HE in situ method, for some false positive patients, and the mismatch patient between primary CY method and SRC method also were discussed in **Supplementary Fig. S12** and **Fig. S13**. In **Supplementary table S5**, it demonstrated that false positive (8/80) of SRC method or false negative (5/80) of SRC method, comparing with PM results. In situ CY&SRC result (2/80) confirms there is no tumor cell of our detection cells when SRC mismatched with PM to get false negative. False negative also may be induced by the bias of sample random collection of ascites. If the high throughput SRC cell detection enables in future, we may improve this condition. Moreover, the PM positive rate of gastric patients decreased significantly after chemotherapy (**Supplementary Fig. S14**), and positive probability of 3 PM+ patient decreased from 85% to 38%; 2 PM- patient decreased from 38% to 27%. This indicates that our SRC method may effectively assess chemotherapy prognosis.

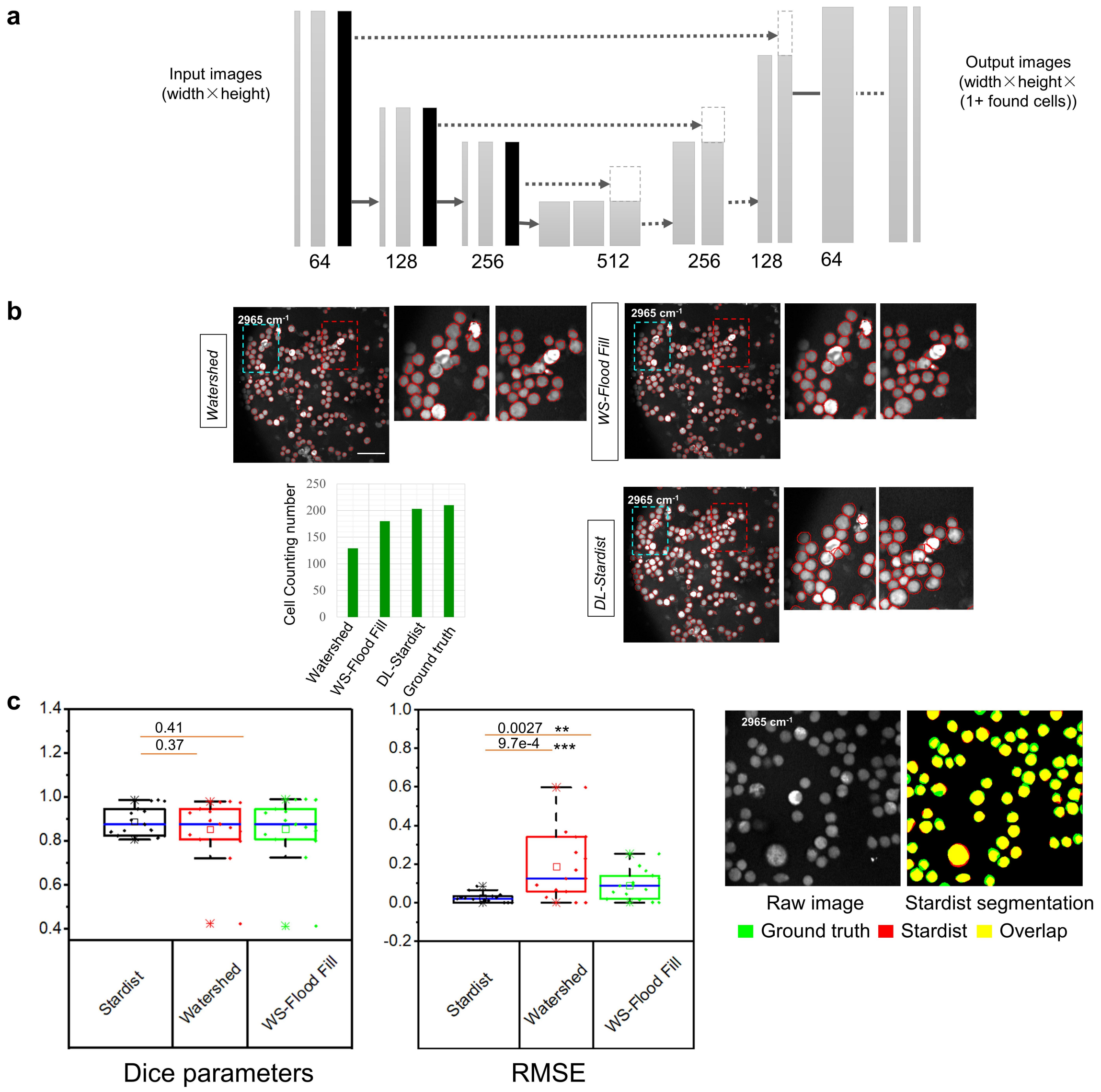


**Fig. S1 | Performance of single cell segmentation by *stardist* model. a**, Neural networks of *stardist* model. **b**, Typical segmentation results of watershed (WS) combing with fill flood method by using *imagej*, watershed by using standard *opencv* library, deep learning (DL) based *stardist* segmentation model. Cell count using watershed; WS-Flood Fill method; stardist; ground truth by manual visual judgment. Red line label the contours of cells, and two zoom-in regions. **c**, Dice parameter and relative root squared error (RRSE) between automated segmentation and ground truth (N=30), and cell segmentation overlay comparison between visual judgment and *stardist*. The box and whisker plots represent median values (center lines), mean values (horizontal bars), minimum and maximum (outliers), 25th to 75th percentiles (box edges) and 1.5x interquartile range (whiskers), with all points plotted. ***<0.0005, **<0.005, *<0.05, Scale bar: 20μm.

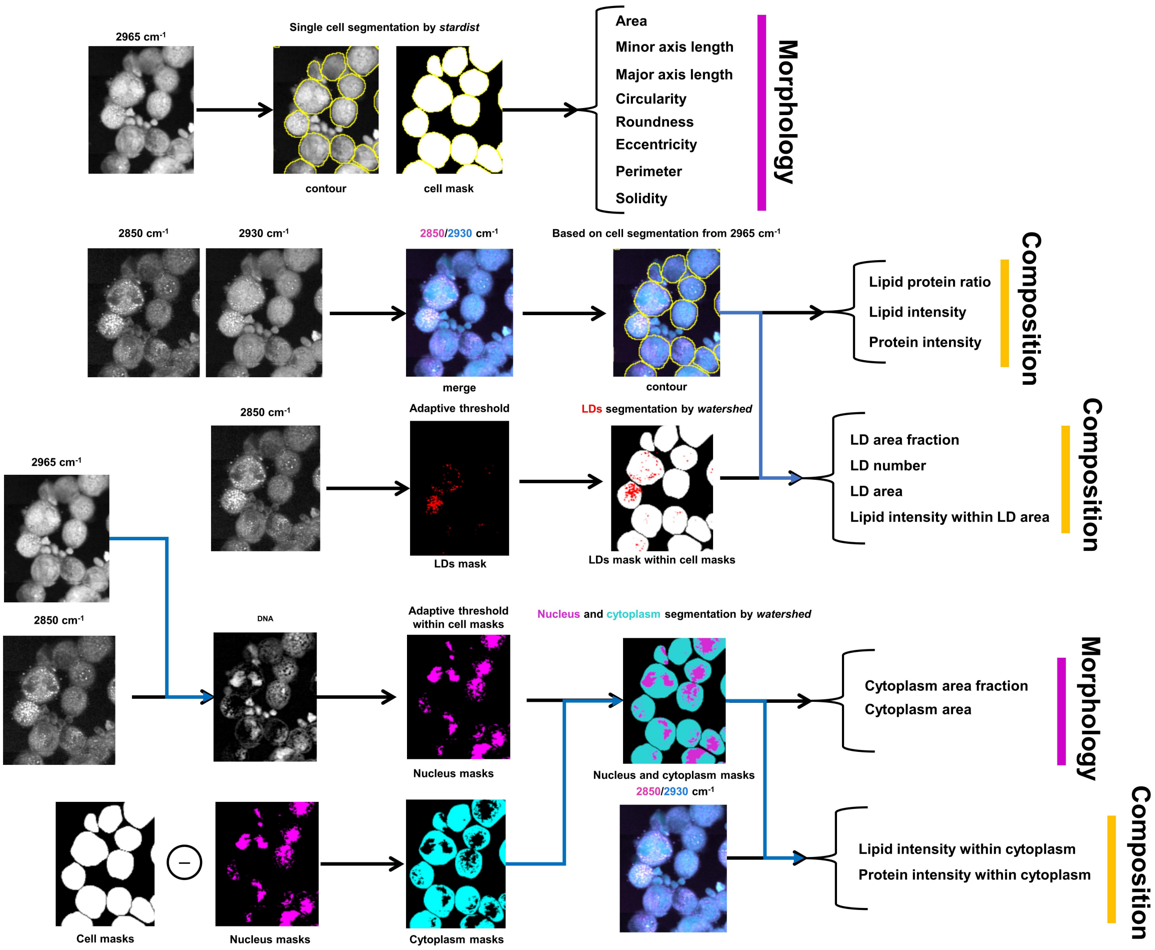


**Fig. S2 | Workflow of feature extraction based on masks.** Firstly, 8 morphology features extract from cell segmentation based on 2965cm$^{-1}$ channel, and 3 composition features extract from 2850cm$^{-1}$ and 2930cm$^{-1}$ based on single cell segmentation mask. Finally, 6 composition features and 2 morphology features extracted from 2850cm$^{-1}$ and 2930cm$^{-1}$ based on single cell mask and LDs, nucleus and cytoplasm masks.

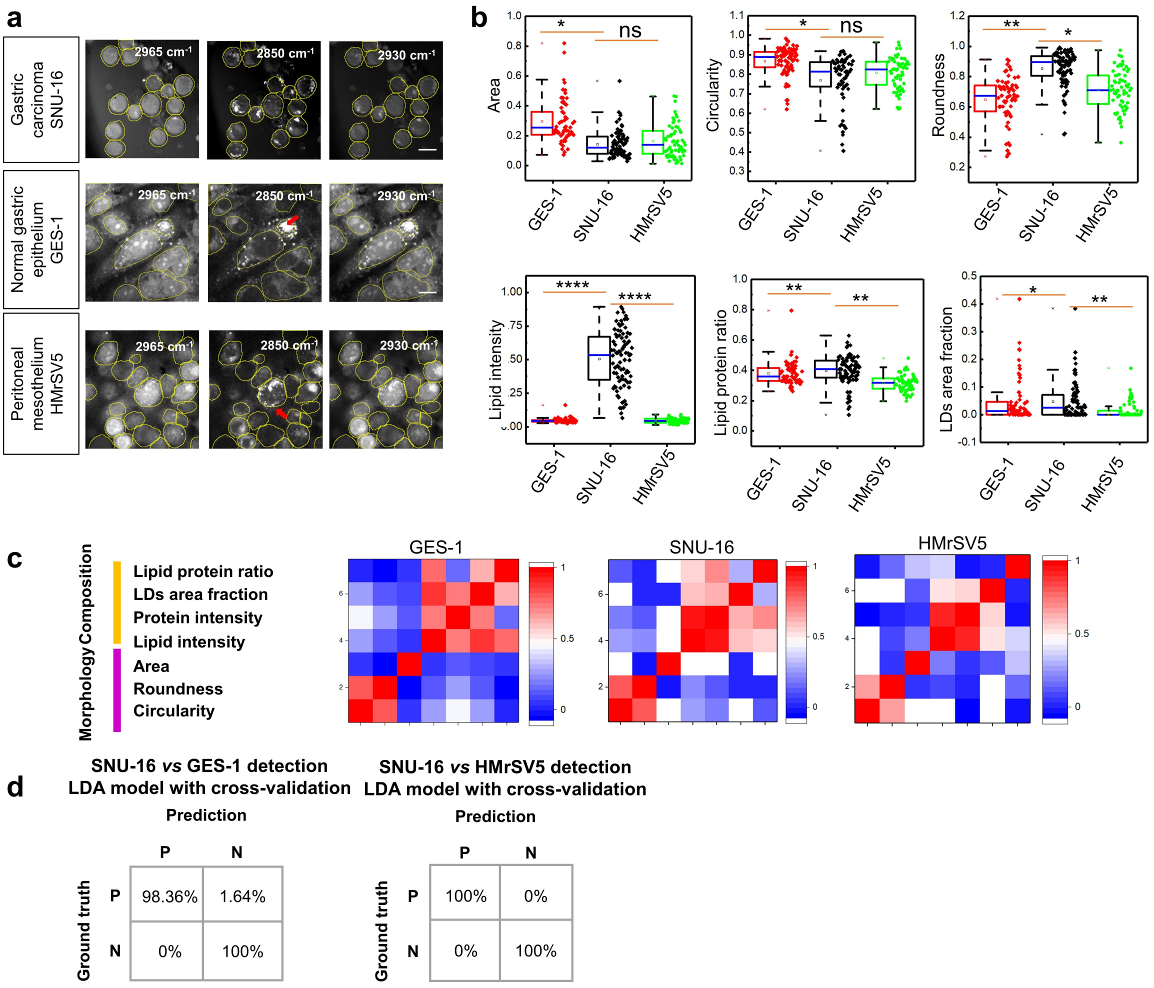


**Fig. S3 | Results of SRC of gastric cell lines with single cell segmentation and classification. a**, Typical three-color SRS images of GES-1 (gastric epithelium cells), SNU-16 (differentiated carcinoma cells), and HMrSV5 (mesothelial cells). **b**, Three morphology features (area, roundness and circularity) and four composition features (lipid intensity, lipid protein ratio, LDs area fraction) comparisons of single cell between gastric cells. The box and whisker plots represent median values (center lines), mean values (horizontal bars), minimum and maximum (outliers), 25th to 75th percentiles (box edges) and 1.5x interquartile range (whiskers), with all points plotted. **c**, Correlation coefficient mapping about features with each other for gastric cells, **d**, confusion matrix of tumor cell detection by linear discriminate analysis (LDA) model. Scale bar: 20μm. ***<0.0005, **<0.005, *<0.05, ns: no significant difference.

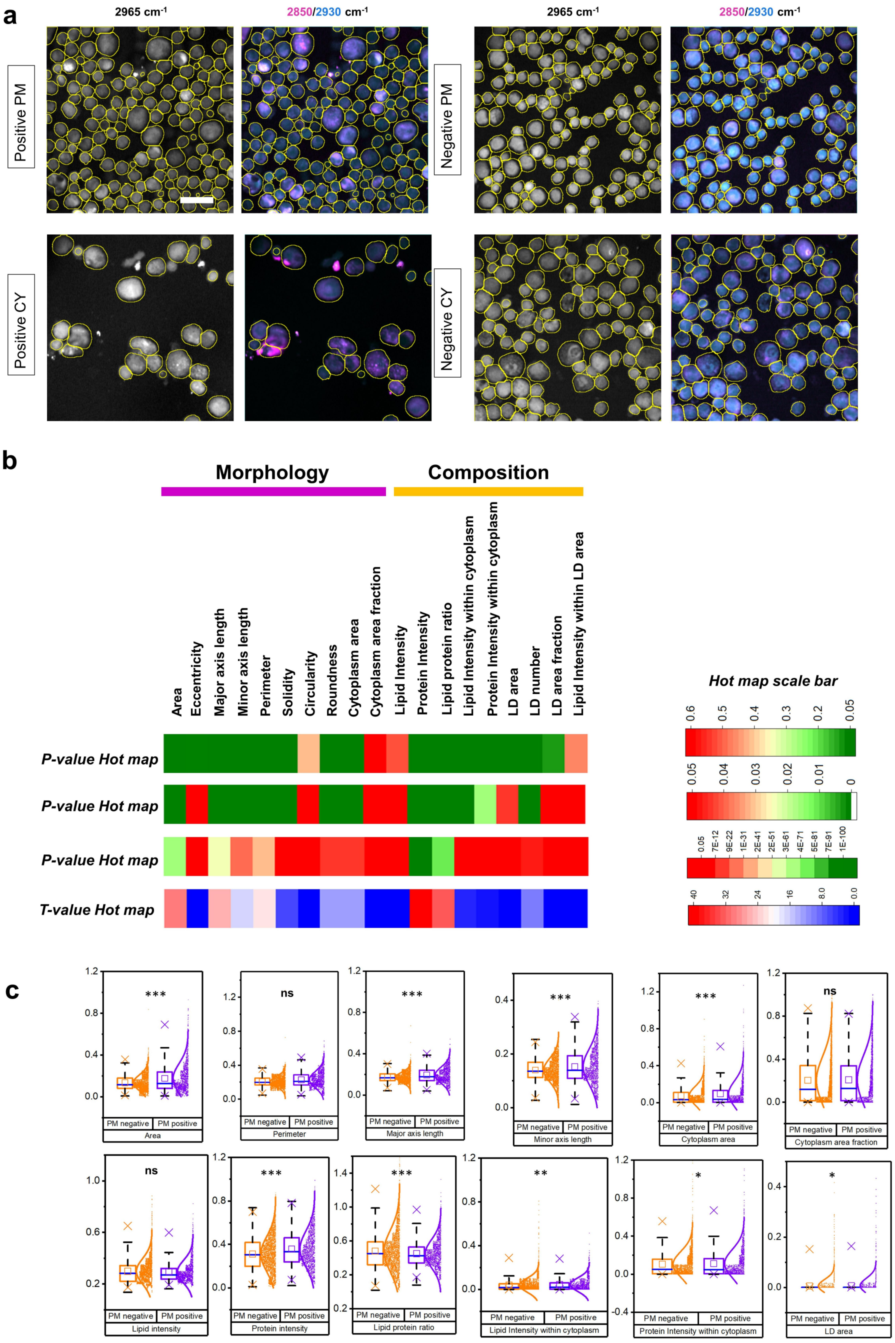


**Fig. S4 | Representative SRS imaging of GC exfoliated cells and comparisons of 19 features between positive and negative PM. a**, Typical images of PM positive and negative specimen. **b**, Hot map of p-values, and t-value by comparing PM positive and negative raw features with three different scale-bar, the features mainly have three types of morphology, composition, and lipid droplets (LD). **c**, feature comparisons between negative and positive PM. scale bar: 20μm. The box and whisker plots represent median values (center lines), mean values (horizontal bars), minimum and maximum (outliers), 25th to 75th percentiles (box edges) and 1.5x interquartile range (whiskers), with all points plotted. ***<0.0005, **<0.005, *<0.05, ns: no significant difference.

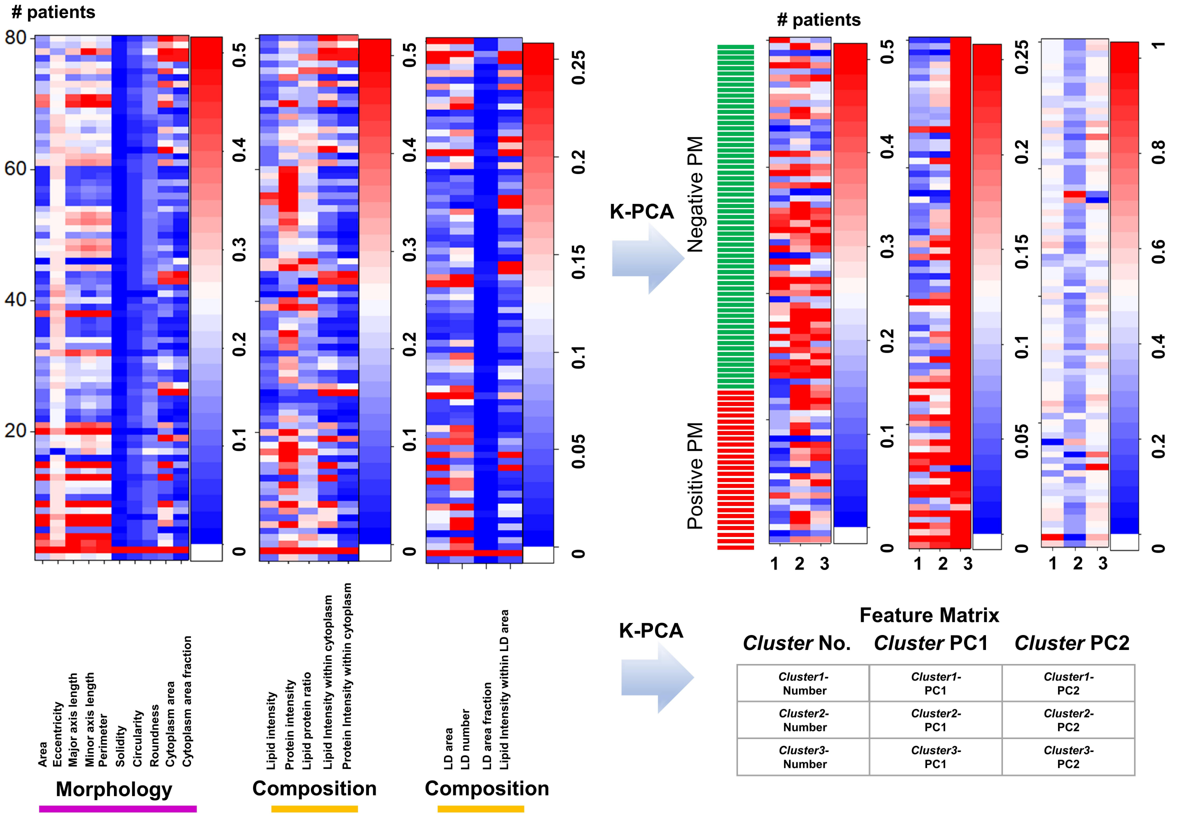


**Fig. S5 | Hot map of features raw feature and feature matrix transformed by K-PCA. a**, 0-1 normalization of average values of raw features before K-PCA. **b**, 0-1 normalization of feature matrix after K-PCA based dimensional reduction. *Cluster1*-number and *Clutser1*-PC1 were obviously different between PM positive and negative.

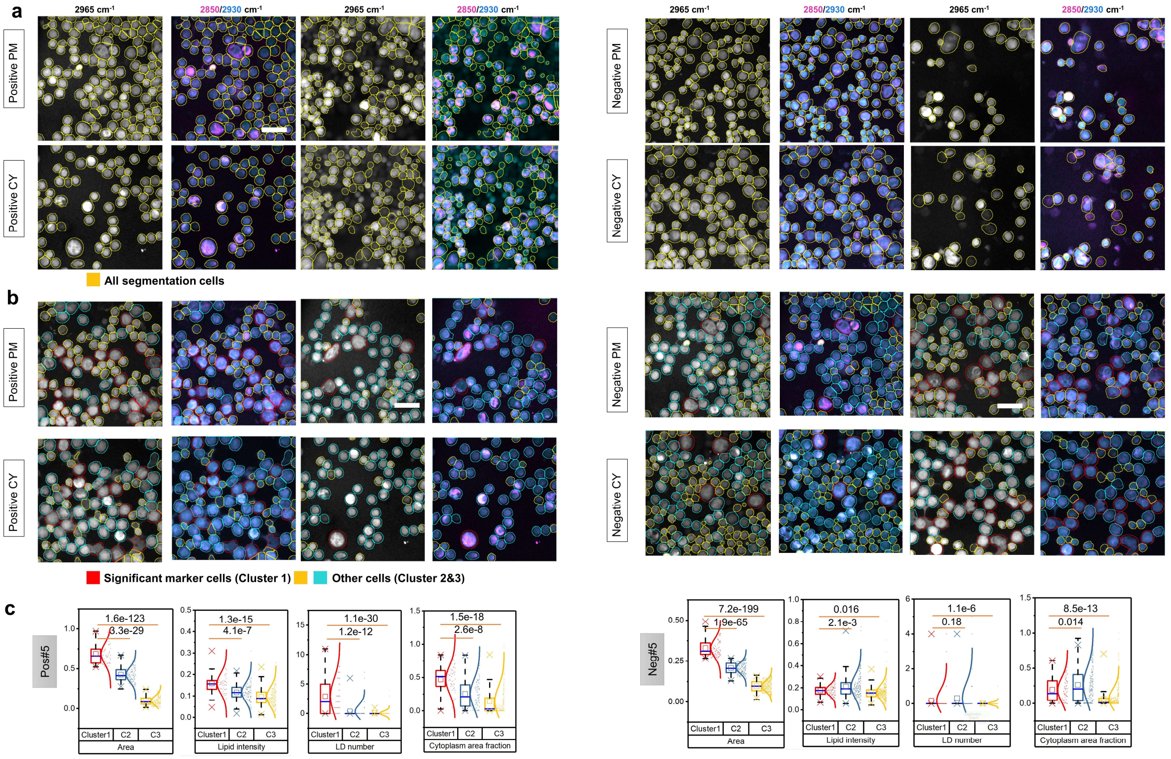


**Fig. S6 | Representative images and quantitative analysis of exfoliated cell phenotyping. a**, Typical cell segmentation results. Yellow labels the all contours. **b**, Typical cell phenotyping results by K-PCA. Red labels *Cluster 1*, cyan labels *Cluster 2* and yellow labels *Cluster 3*. **c**, Features (LD number, lipid intensity, cellular area and cytoplasm area fraction) quantification and comparisons among cell clusters in CY NEG/POS#5. The numbers with underlines denote $p$ values. The box and whisker plots represent median values (center lines), mean values (horizontal bars), minimum and maximum (outliers), 25th to 75th percentiles (box edges) and 1.5x interquartile range (whiskers), with all points plotted.

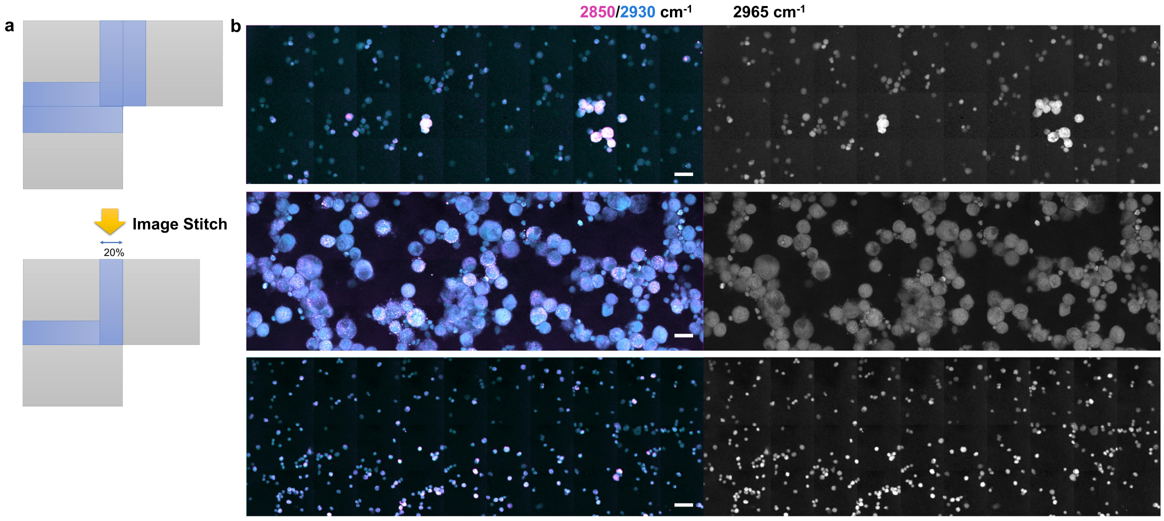


**Fig. S7 | Image stitching of three-color SRS imaging of GC exfoliated cells. a**, Schematic of image stitching, image tiles generated using the 20% overlap approach are stitched to an absolute Cartesian coordinate system. **b**, Typical three-color SRS imaging of GC exfoliated cells after image stitching. Upper shows typical cytology positive specimen with low percentage of tumor cells, and medium shows typical cytology specimen with high percentage of tumor cells, and lower shows typical cytology negative specimen without tumor cells. Scale bar: 20μm.

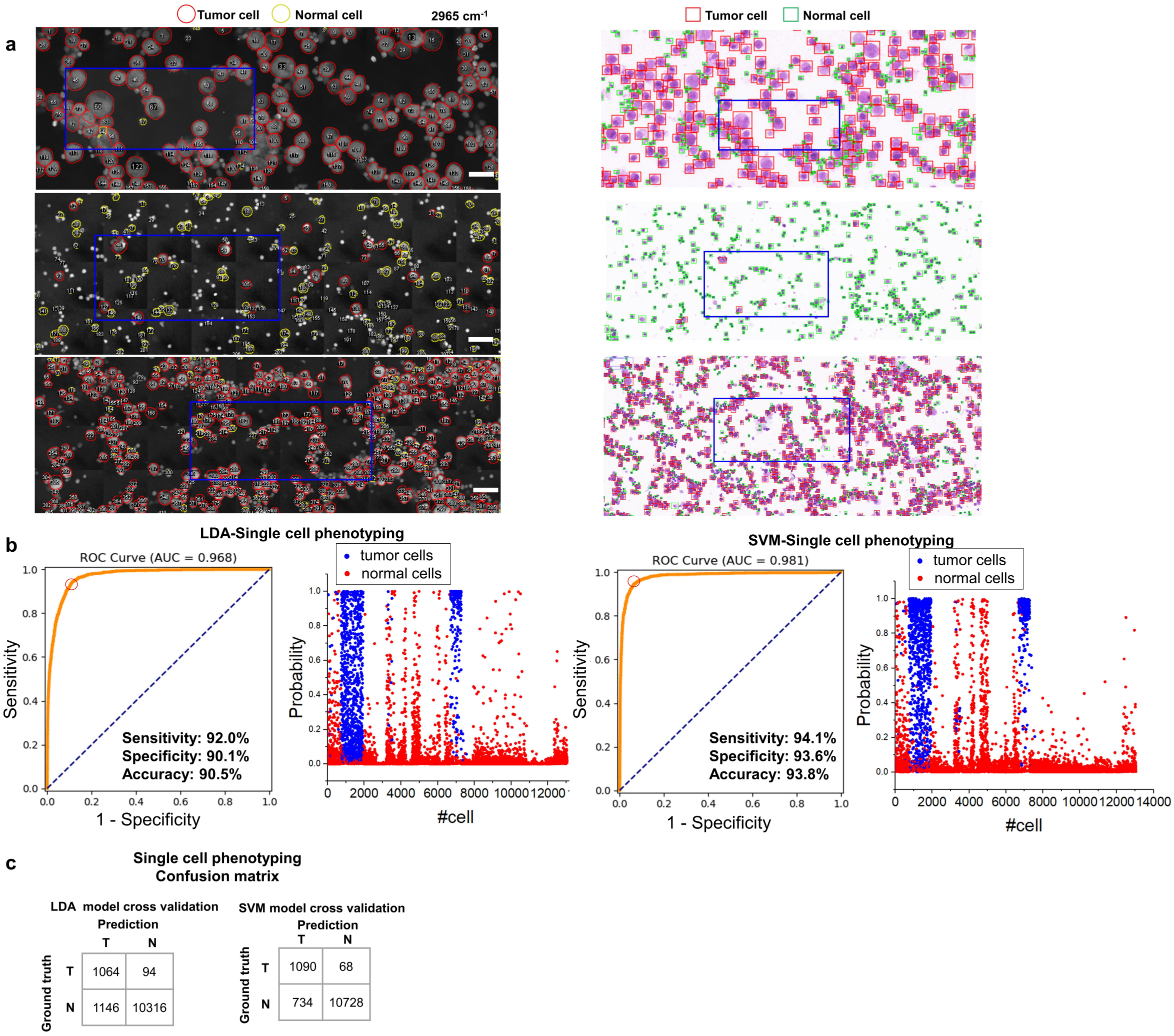


**Fig. S8 | In situ SRS&HE imaging of GC exfoliated cells.** Results of single cell phenotyping by machine learning (ML) classifiers (SVM and LDA). **a**, Typical H&E&SRS images of GC cells with cell phenotyping results. We collected imaging data to create the training and test dataset include GC exfoliated cells (1158 normal cells; 11462 tumor cells). **b**, ROC curve (left), the AUCs, sensitivity, specificity and accuracy and the positive probability of cell phenotyping (right), using ML algorithms. **c**, Confusion matric of single cell phenotyping using ML algorithms.

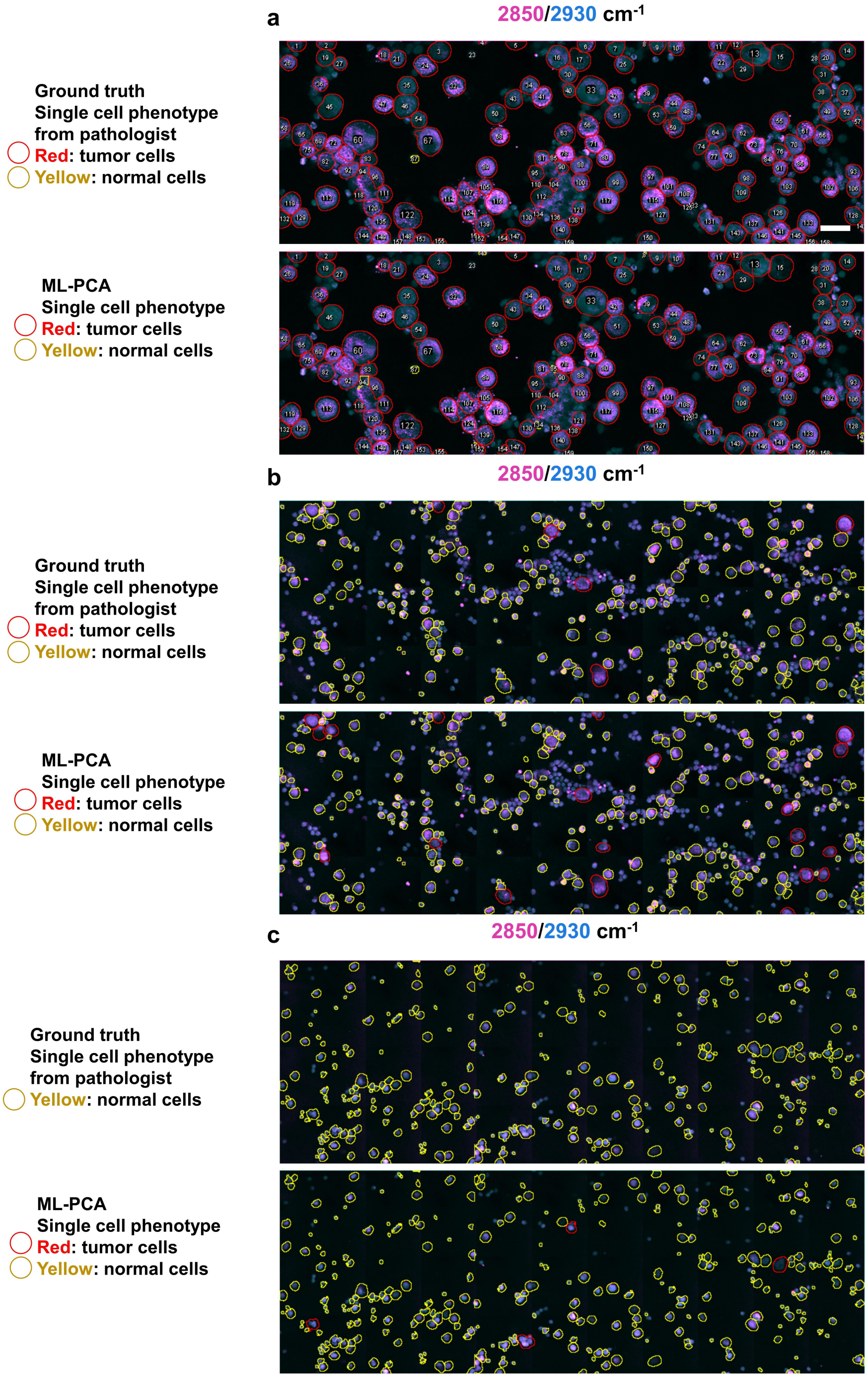


**Fig. S9 | Typical tumor cell and normal cell detection by ML-PCA.** Comparisons of ML-PCA results and H&E results for three types of specimens. SRS&HE in situ images were labeled by two senior pathologists. **a**, Typical PM positive specimen with high percentage of tumor cells. **b**, Typical PM positive specimen with low percentage of tumor cells. **c**, Typical PM negative specimen without tumor cells.

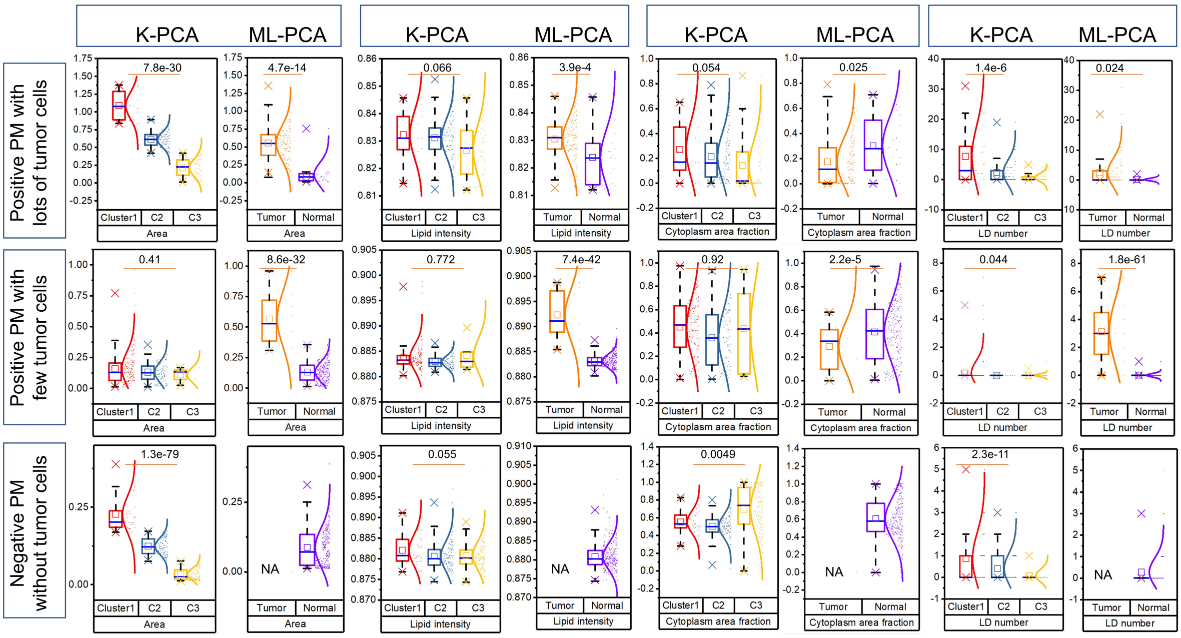


**Fig. S10 | Results of feature profiles by K-PCA and ML-PCA.** We compared the features including area, lipid intensity, LD number, cytoplasm area fraction of three typical specimen with p-value. PM positive specimen with high percentage of tumor cells. Typical PM positive specimen with low percentage of tumor cells. Typical PM negative specimen without tumor cells. The box and whisker plots represent median values (center lines), mean values (horizontal bars), minimum and maximum (outliers), 25th to 75th percentiles (box edges) and 1.5x interquartile range (whiskers), with all points plotted.

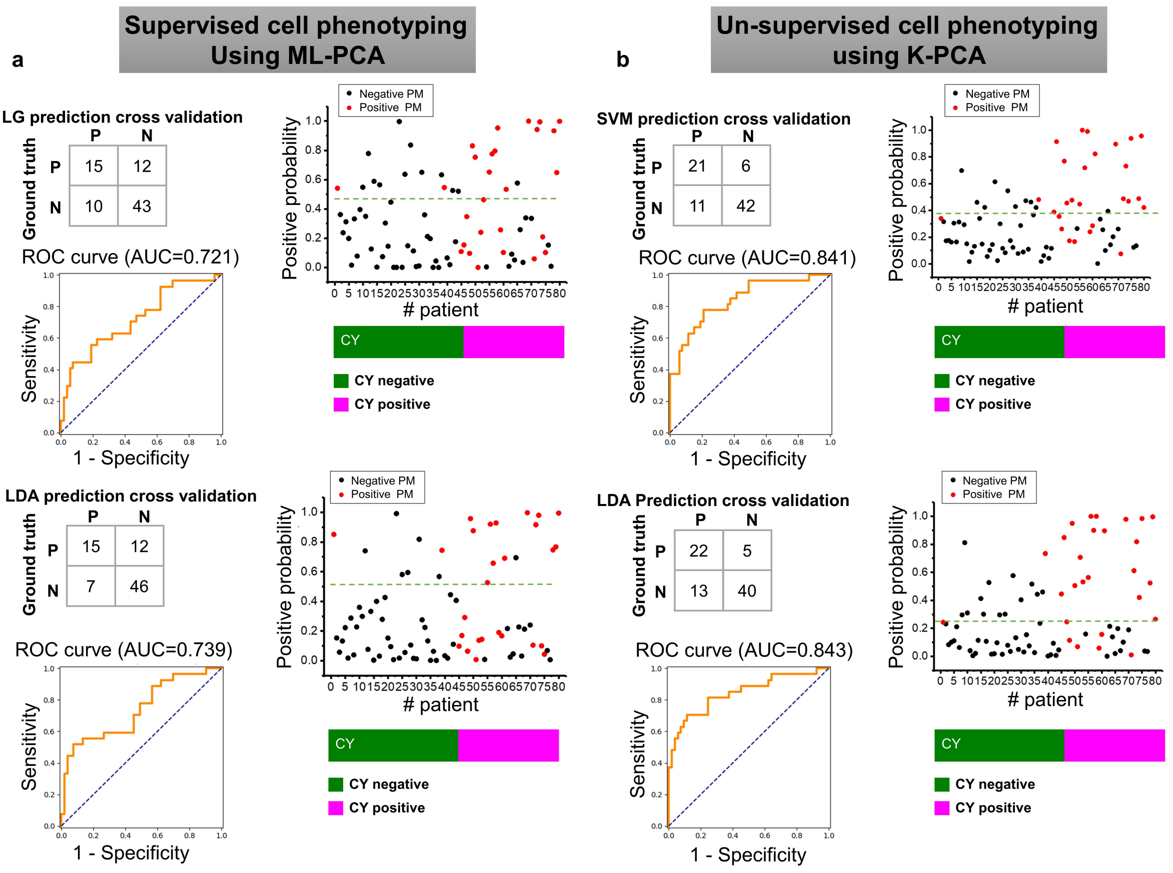


**Fig. S11 | Results of PM detection by classifiers (SVM, LDA, LG etc.).** The input feature matrix respectively comes from supervised ML-PCA and unsupervised K-PCA cell phenotyping methods. PM positive probability with cross-validation, confusion matrix and ROC curves of 80 patients (27 positive PM; 53 negative PM, 35 positive cytology; 45 negative cytology). **a**, Using ML-PCA algorithm, the AUCs were respectively 0.797, 0.739 and 0.721 by SVM, LDA, and LG. The best performance of ML-PCA and SVM model was 0.797 described in **Fig. 4**, ML-PCA and LG/LDA was here. **b**, Using K-PCA algorithm, the AUCs were respectively 0.841, 0.85 and 0.843 by SVM, LDA, and LG. The best performance of K-PCA and LG was 0.85 described in **Fig. 4**, K-PCA and SVM/LDA was here.

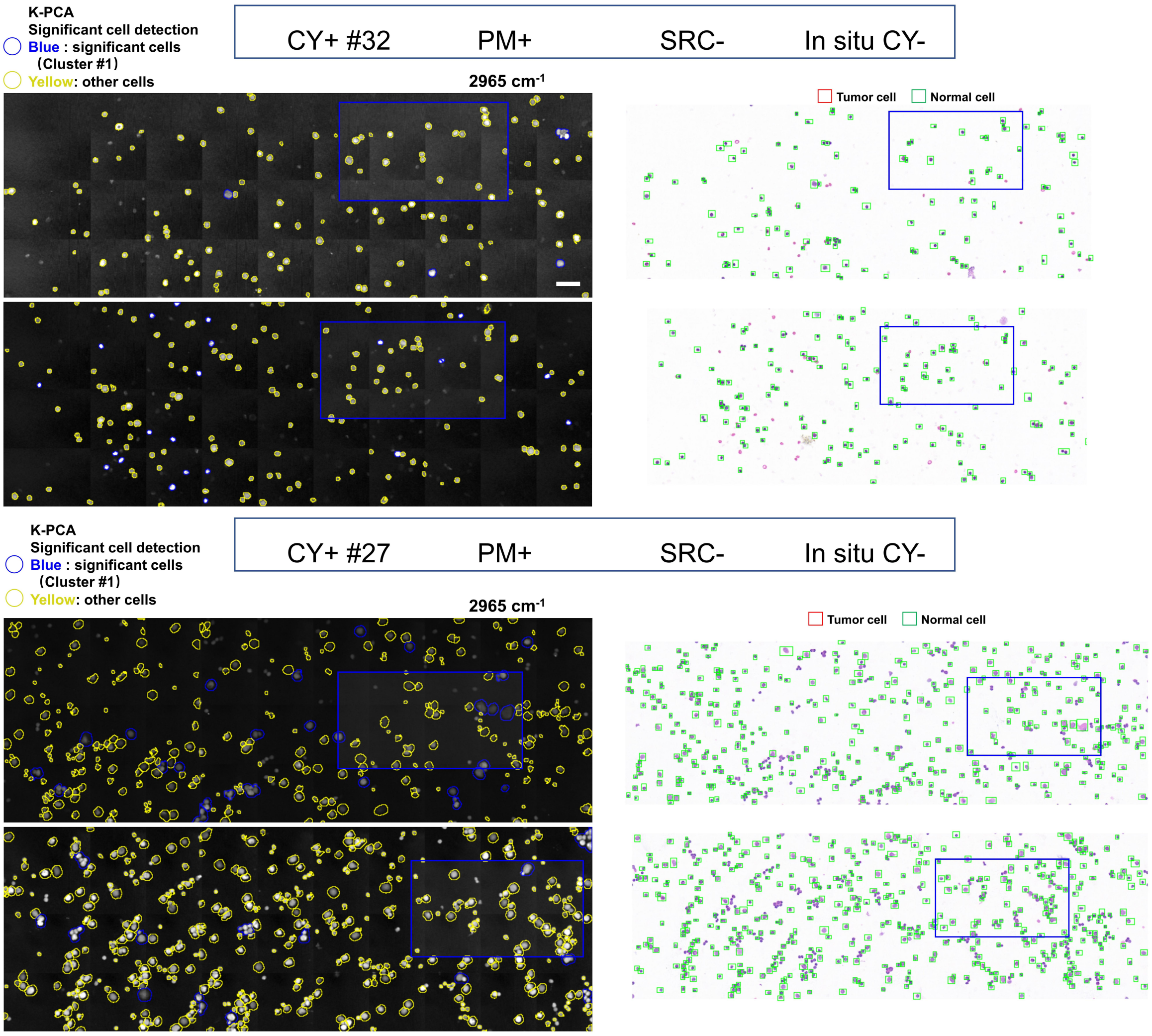


**Fig. S12 | Typical SRS&HE in-situ images for SRC negative detection.** The mismatch between CY+, PM+ and our SRC- method.

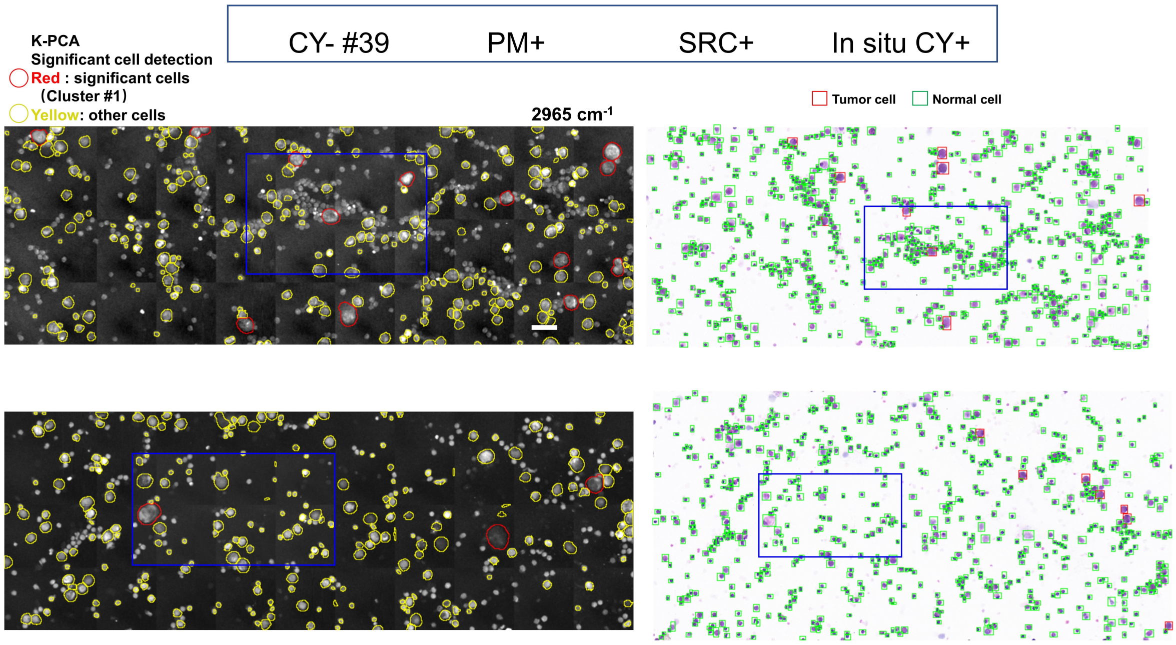


**Fig. S13 | Typical SRS&HE in-situ images for SRC positive detection.** The mismatch between CY-, PM+ and our SRC+ method.

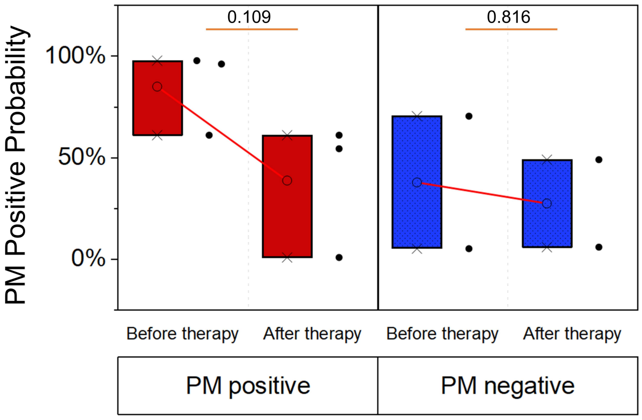


**Fig. S14 | PM positive probability before and after chemotherapy for patients diagnosed with PM positive (n = 3) and PM negative (n=2).** The numbers with underlines denote *p* values. The box and whisker plots represent median values (center lines), mean values (horizontal bars), minimum and maximum (outliers), 25th to 75th percentiles (box edges) and 1.5x interquartile range (whiskers), with all points plotted.

**Table S1 | Typical raw features with 0-1 normalization from 80 patients.**

| Patient ID | Conventional Cytology results | Histopathology results | Area | Lipid intensity | Protein intensity | Cytoplasm area fraction | LD number |
|---|---|---|---|---|---|---|---|
| NEG#1* | negative | positive | 0.192 | 0.114 | 0.285 | 0.027 | 0.073 |
| NEG#2 | negative | negative | 0.488 | 1 | 0.947 | 1 | 1 |
| NEG#3 | negative | negative | 0.385 | 0.165 | 0.387 | 0.066 | 0.206 |
| NEG#4 | negative | negative | 0.460 | 0.171 | 0.251 | 0.047 | 0.306 |
| NEG#5 | negative | negative | 0.049 | 0.072 | 0.177 | 0.004 | 0.014 |
| NEG#6 | negative | negative | 0.619 | 0.316 | 0.694 | 0.084 | 0.263 |
| NEG#7 | negative | negative | 0.774 | 0.083 | 0.095 | 0.010 | 0.410 |
| NEG#8 | negative | negative | 0.158 | 0.1116 | 0.357 | 0.181 | 0.086 |
| NEG#9 | negative | negative | 0.694 | 0.326 | 0.747 | 0.125 | 0.441 |
| NEG#10 | negative | negative | 0.056 | 0.093 | 0.212 | 0.008 | 0.042 |
| NEG#11 | negative | negative | 0.155 | 0.152 | 0.364 | 0.028 | 0.109 |
| NEG#12 | negative | negative | 0.164 | 0.167 | 0.469 | 0.048 | 0.194 |
| NEG#13 | negative | negative | 1 | 0.045 | 0.054 | 0.064 | 0.192 |
| NEG#14 | negative | negative | 0.151 | 0.158 | 0.456 | 0.018 | 0.143 |
| NEG#15 | negative | negative | 0.877 | 0.160 | 0.395 | 0.157 | 0.089 |
| NEG#16 | negative | negative | 0.025 | 0.251 | 0.706 | 0.030 | 0.044 |
| NEG#17 | negative | negative | 0.157 | 0.392 | 0.664 | 0.170 | 0.178 |
| NEG#18 | negative | negative | 0.192 | 0.242 | 0.588 | 0.058 | 0.073 |
| NEG#19 | negative | negative | 0.175 | 0.098 | 0.304 | 0.376 | 0.221 |
| NEG#20 | negative | negative | 0.499 | 0.313 | 0.700 | 0.002 | 0.206 |
| NEG#21 | negative | negative | 0.375 | 0.076 | 0.196 | 0.084 | 0.218 |
| NEG#22 | negative | negative | 0.039 | 0.127 | 0.320 | 0.021 | 0.047 |
| NEG#23 | negative | negative | 0.069 | 0.067 | 0.213 | 0.007 | 0.040 |
| NEG#24 | negative | negative | 0.118 | 0.054 | 0.141 | 0.025 | 0.030 |
| NEG#25 | negative | negative | 0.109 | 0.021 | 0.060 | 0.039 | 0.124 |
| NEG#26 | negative | negative | 0.114 | 0.015 | 0.098 | 0.531 | 0.350 |
| NEG#27 | negative | negative | 0.151 | 0.096 | 0.146 | 0.238 | 0.222 |
| NEG#28 | negative | negative | 0.169 | 0.116 | 0.363 | 0.044 | 0.051 |
| NEG#29 | negative | negative | 0.125 | 0.083 | 0.269 | 0.041 | 0.060 |
| NEG#30 | negative | negative | 0.152 | 0.119 | 0.349 | 0.174 | 0.111 |
| NEG#31 | negative | negative | 0.148 | 0.093 | 0.325 | 0.150 | 0.152 |
| NEG#32 | negative | negative | 0.283 | 0.049 | 0.076 | 0.037 | 0.202 |
| NEG#33 | negative | negative | 0.093 | 0.107 | 0.381 | 0 | 0.026 |
| NEG#34 | negative | negative | 0.181 | 0.126 | 0.415 | 0.067 | 0.142 |
| NEG#35 | negative | negative | 0.048 | 0.164 | 0.500 | 0.008 | 0.041 |
| NEG#36 | negative | negative | 0.037 | 0.096 | 0.264 | 0.033 | 0.025 |
| NEG#37 | negative | negative | 0.038 | 0.099 | 0.215 | 0.023 | 0.030 |
| NEG#38 | negative | negative | 0.430 | 0.104 | 0.142 | 0.033 | 0.037 |
| NEG#39* | negative | positive | 0.073 | 0.328 | 0.495 | 0.080 | 0.067 |
| NEG#40 | negative | negative | 0.133 | 0.140 | 0.417 | 0.028 | 0.060 |
| NEG#41 | negative | negative | 0.066 | 0 | 0 | 0.132 | 0.130 |
| NEG#42 | negative | negative | 0.102 | 0.071 | 0.146 | 0.040 | 0.098 |
| NEG#43 | negative | negative | 0.066 | 0.021 | 0.156 | 0.455 | 0.296 |
| NEG#44 | negative | negative | 0.04 | 0.077 | 0.161 | 0.484 | 0.402 |
| NEG#45 | negative | negative | 0.215 | 0.283 | 0.507 | 0.103 | 0.046 |
| POS#1 | positive | positive | 0 | 0.284 | 0.337 | 0.250 | 0.105 |
| POS#2 | positive | positive | 0.222 | 0.098 | 0.282 | 0.013 | 0.129 |
| POS#3 | positive | positive | 0.260 | 0.109 | 0.218 | 0.059 | 0.139 |
| POS#4 | positive | positive | 0.227 | 0.136 | 0.274 | 0.018 | 0.02 |

| | | | | | | | |
|---|---|---|---|---|---|---|---|
| POS#5 | positive | positive | 0.091 | 0.107 | 0.315 | 0.008 | 0.040 |
| POS#6 | positive | positive | 0.190 | 0.142 | 0.378 | 0.025 | 0.091 |
| POS#7 | positive | positive | 0.256 | 0.196 | 0.436 | 0.022 | 0.077 |
| POS#8 | positive | positive | 0.231 | 0.170 | 0.379 | 0.033 | 0.120 |
| POS#9 | positive | positive | 0.173 | 0.180 | 0.508 | 0.141 | 0.186 |
| POS#10* | positive | negative | 0.045 | 0.384 | 1 | 0.074 | 0 |
| POS#11 | positive | positive | 0.063 | 0.465 | 0.975 | 0.133 | 0.029 |
| POS#12 | positive | positive | 0.067 | 0.171 | 0.511 | 0.064 | 0.057 |
| POS#13 | positive | positive | 0.048 | 0.202 | 0.550 | 0.168 | 0.044 |
| POS#14 | positive | positive | 0.041 | 0.273 | 0.795 | 0.035 | 0.061 |
| POS#15 | positive | positive | 0.036 | 0.214 | 0.470 | 0.028 | 0 |
| POS#16 | positive | positive | 0.289 | 0.115 | 0.161 | 0.168 | 0.157 |
| POS#17 | positive | positive | 0.218 | 0.115 | 0.269 | 0.011 | 0.128 |
| POS#18* | positive | negative | 0.175 | 0.109 | 0.184 | 0.282 | 0.248 |
| POS#19* | positive | negative | 0.143 | 0.165 | 0.313 | 0.178 | 0.085 |
| POS#20* | positive | negative | 0.185 | 0.174 | 0.256 | 0.086 | 0.099 |
| POS#21* | positive | negative | 0.133 | 0.128 | 0.270 | 0.218 | 0.177 |
| POS#22* | positive | negative | 0.144 | 0.087 | 0.154 | 0.053 | 0.034 |
| POS#23* | positive | negative | 0.175 | 0.126 | 0.348 | 0.056 | 0.068 |
| POS#24* | positive | negative | 0.135 | 0.052 | 0.155 | 0.014 | 0.029 |
| POS#25 | positive | positive | 0.394 | 0.182 | 0.528 | 0.142 | 0.566 |
| POS#26* | positive | negative | 0.367 | 0.076 | 0.190 | 0.106 | 0.189 |
| POS#27 | positive | positive | 0.191 | 0.084 | 0.200 | 0.024 | 0.105 |
| POS#28 | positive | positive | 0.190 | 0.079 | 0.104 | 0.294 | 0.122 |
| POS#29 | positive | positive | 0.088 | 0.129 | 0.395 | 0.021 | 0.031 |
| POS#30 | positive | positive | 0.217 | 0.187 | 0.249 | 0.234 | 0.162 |
| POS#31 | positive | positive | 0.180 | 0.091 | 0.166 | 0.085 | 0.102 |
| POS#32* | positive | negative | 0.076 | 0.188 | 0.222 | 0.483 | 0.135 |
| POS#33 | positive | positive | 0.290 | 0.081 | 0.191 | 0.478 | 0.665 |
| POS#34 | positive | positive | 0.141 | 0.101 | 0.279 | 0.265 | 0.087 |
| POS#35 | positive | positive | 0.157 | 0.076 | 0.125 | 0.405 | 0.437 |

* represents the mismatch between cytological and histopathological results.

**Table S2 | Values of *Cluster1*-PC1, *Cluster1*-PC2 and *Cluster2*-PC1, *Cluster2*-PC2 of 80 patients.**

| Patient ID | Cytological results | Laparoscopic results | Cluster1-PC1 | Cluster2-PC1 | Cluster1-PC2 | Cluster2-PC2 |
|---|---|---|---|---|---|---|
| NEG#1* | negative | positive | 1737.094 | 408.314 | 173.373 | -76.369 |
| NEG#2 | negative | negative | 844.614 | 89.835 | -7.123 | -26.401 |
| NEG#3 | negative | negative | 445.168 | -117.110 | -24.275 | 19.424 |
| NEG#4 | negative | negative | 938.398 | 150.506 | -10.108 | -4.268 |
| NEG#5 | negative | negative | 708.001 | 227.756 | -59.817 | 60.633 |
| NEG#6 | negative | negative | 1158.821 | 180.735 | -15.841 | -20.370 |
| NEG#7 | negative | negative | 1027.32 | 215.410 | 27.431 | -43.363 |
| NEG#8 | negative | negative | 957.964 | 240.060 | -117.903 | 70.044 |
| NEG#9 | negative | negative | 902.757 | 84.886 | 102.554 | -16.743 |
| NEG#10 | negative | negative | 1225.35 | 424.159 | -103.639 | 80.883 |
| NEG#11 | negative | negative | 757.296 | 101.472 | 56.199 | -46.566 |
| NEG#12 | negative | negative | 457.147 | 27.520 | 25.156 | -24.349 |
| NEG#13 | negative | negative | 797.009 | 143.337 | -1.691 | -22.385 |
| NEG#14 | negative | negative | 666.129 | 180.469 | -76.770 | 51.629 |
| NEG#15 | negative | negative | 260.560 | -80.291 | -24.717 | 670.329 |
| NEG#16 | negative | negative | 1284.856 | -46.309 | -37.757 | 381.138 |
| NEG#17 | negative | negative | 715.981 | 237.612 | 94.674 | -30.277 |
| NEG#18 | negative | negative | 729.73 | 227.736 | 7.153 | 1.716 |
| NEG#19 | negative | negative | 821.525 | 252.443 | 100.075 | -79.735 |
| NEG#20 | negative | negative | 478.623 | 111.957 | -75.776 | 50.604 |
| NEG#21 | negative | negative | 974.253 | 353.495 | 104.500 | -40.222 |
| NEG#22 | negative | negative | 1520.546 | 561.952 | 208.092 | -98.595 |
| NEG#23 | negative | negative | 816.703 | 213.091 | 54.045 | -58.634 |
| NEG#24 | negative | negative | 1777.592 | 720.063 | -280.841 | 89.608 |
| NEG#25 | negative | negative | 481.863 | 96.590 | -152.683 | 133.818 |
| NEG#26 | negative | negative | 739.063 | 330.870 | 110.313 | -107.104 |
| NEG#27 | negative | negative | 2378.29 | 435.110 | -97.483 | 1.340 |
| NEG#28 | negative | negative | 823.981 | 231.311 | -84.849 | 84.967 |
| NEG#29 | negative | negative | 664.221 | 9.947 | 54.300 | -50.059 |
| NEG#30 | negative | negative | 1320.432 | 488.849 | -61.390 | -6.251 |
| NEG#31 | negative | negative | 732.102 | -20.216 | -49.950 | 84.793 |
| NEG#32 | negative | negative | 920.558 | 61.997 | 113.268 | -54.742 |
| NEG#33 | negative | negative | 763.668 | 268.196 | -26.423 | 32.675 |
| NEG#34 | negative | negative | 1126.836 | 361.082 | 26.231 | -9.368 |
| NEG#35 | negative | negative | 533.805 | 113.539 | 6.119 | 0.049 |
| NEG#36 | negative | negative | 870.815 | 91.880 | -66.679 | 42.088 |
| NEG#37 | negative | negative | 904.256 | 144.142 | 36.072 | -24.010 |
| NEG#38 | negative | negative | 1301.845 | 173.017 | 46.695 | -57.655 |
| NEG#39* | negative | positive | 1509.39 | 308.400 | -35.687 | 28.320 |
| NEG#40 | negative | negative | 476.672 | -60.531 | 2.140 | -28.408 |
| NEG#41 | negative | negative | 685.154 | 23.928 | -69.651 | 161.737 |
| NEG#42 | negative | negative | 692.580 | 131.260 | 72.172 | -59.008 |
| NEG#43 | negative | negative | 518.024 | -64.426 | -6.299 | 11.182 |
| NEG#44 | negative | negative | 870.637 | 128.515 | -21.545 | 16.332 |
| NEG#45 | negative | negative | 732.973 | 211.145 | 48.172 | -35.386 |
| POS#1 | positive | positive | 1351.002 | 217.660 | 45.233 | -12.268 |
| POS#2 | positive | positive | 2236.688 | 158.635 | 75.074 | -51.025 |
| POS#3 | positive | positive | 965.951 | 230.955 | 203.012 | -68.181 |
| POS#4 | positive | positive | 893.065 | 174.681 | 123.538 | -30.362 |

| | | | | | | |
|---|---|---|---|---|---|---|
| POS#5 | positive | positive | 1950.394 | 580.662 | -257.156 | 195.820 |
| POS#6 | positive | positive | 1514.125 | 343.351 | 24.778 | -18.591 |
| POS#7 | positive | positive | 808.706 | 15.660 | 5.539 | -6.503 |
| POS#8 | positive | positive | 1129.354 | 114.743 | -92.609 | 22.080 |
| POS#9 | positive | positive | 1631.954 | 781.733 | -833.677 | 312.455 |
| POS#10* | positive | negative | 733.400 | 70.399 | 0.751 | -17.881 |
| POS#11 | positive | positive | 1211.185 | 248.607 | -116.690 | 54.805 |
| POS#12 | positive | positive | 6433.348 | 1047.681 | 50.451 | -315.115 |
| POS#13 | positive | positive | 1942.708 | 557.735 | -253.926 | 62.929 |
| POS#14 | positive | positive | 3089.619 | 46.630 | 58.197 | -144.437 |
| POS#15 | positive | positive | 905.032 | 644.500 | -322.444 | 232.145 |
| POS#16 | positive | positive | 1079.778 | 196.359 | 23.834 | -25.271 |
| POS#17 | positive | positive | 2114.345 | 685.251 | -71.387 | 59.997 |
| POS#18* | positive | negative | 531.640 | -83.896 | 29.986 | -74.706 |
| POS#19* | positive | negative | 953.422 | 114.940 | -6.245 | -6.744 |
| POS#20* | positive | negative | 929.428 | 125.100 | -46.843 | 8.561 |
| POS#21* | positive | negative | 827.804 | -67.098 | 99.496 | -139.662 |
| POS#22* | positive | negative | 1980.221 | 594.470 | -99.397 | 7.341 |
| POS#23* | positive | negative | 894.292 | 106.843 | 153.310 | -71.751 |
| POS#24* | positive | negative | 948.777 | 57.226 | -26.856 | 39.538 |
| POS#25 | positive | positive | 2243.022 | 504.56 | -104.258 | 84.040 |
| POS#26* | positive | negative | 929.046 | 238.233 | -95.467 | 138.634 |
| POS#27 | positive | positive | 754.143 | 113.493 | -12.433 | 26.555 |
| POS#28 | positive | positive | 1590.997 | 62.701 | 65.302 | -95.256 |
| POS#29 | positive | positive | 2152.295 | 513.800 | 123.822 | -69.771 |
| POS#30 | positive | positive | 1038.345 | 81.668 | 100.223 | -18.534 |
| POS#31 | positive | positive | 2356.749 | 326.469 | -145.170 | 4.644 |
| POS#32* | positive | negative | 662.761 | 41.833 | 8.090 | -3.524 |
| POS#33 | positive | positive | 1409.766 | 262.881 | 8.388 | -1.547 |
| POS#34 | positive | positive | 8111.398 | 2180.946 | 878.87 | -282.361 |
| POS#35 | positive | positive | 2665.016 | 587.554 | 23.094 | 13.085 |

* represents the mismatch between the cytological and histopathological results.

**Table S3 | Diagnostic results for each patient, including conventional cytology, histopathology, and SRC (based on SVM, LDA, or LG models) predicted probability of PM.**

| Patient ID | Cytological results | Laparoscopic results | age | sex | SRC (SVM) positive probability (%) | SRC (LDA) positive probability (%) | SRC (LG) positive probability (%) |
|---|---|---|---|---|---|---|---|
| NEG#1* | negative | positive | 60 | male | 33.77 | 28.54 | 43.96 |
| NEG#2 | negative | negative | 57 | male | 34.94 | 39.68 | 29.29 |
| NEG#3 | negative | negative | 42 | male | 15.34 | 0.77 | 10.61 |
| NEG#4 | negative | negative | 62 | male | 20.97 | 0.90 | 12.62 |
| NEG#5 | negative | negative | 70 | female | 20.23 | 7.49 | 8.21 |
| NEG#6 | negative | negative | 72 | female | 30.91 | 27.28 | 32.13 |
| NEG#7 | negative | negative | 52 | male | 13.01 | 2.96 | 6.26 |
| NEG#8 | negative | negative | 44 | female | 31.66 | 39.19 | 22.13 |
| NEG#9 | negative | negative | 60 | male | 66.44 | 90.85 | 90.86 |
| NEG#10 | negative | negative | 59 | male | 31.85 | 10.73 | 33.42 |
| NEG#11 | negative | negative | 63 | male | 10.98 | 3.90 | 3.50 |
| NEG#12 | negative | negative | 74 | male | 4.27 | 0.89 | 0.05 |
| NEG#13 | negative | negative | 64 | male | 11.35 | 1.84 | 3.66 |
| NEG#14 | negative | negative | 52 | male | 15.01 | 14.42 | 9.61 |
| NEG#15 | negative | negative | 63 | male | 44.87 | 98.41 | 99.62 |
| NEG#16 | negative | negative | 48 | male | 35.46 | 99.95 | 16.44 |
| NEG#17 | negative | negative | 59 | male | 17.93 | 5.46 | 5.37 |
| NEG#18 | negative | negative | 47 | male | 48.60 | 56.37 | 25.17 |
| NEG#19 | negative | negative | 68 | male | 3.28 | 0.80 | 0.76 |
| NEG#20 | negative | negative | 57 | male | 6.12 | 0.10 | 0.46 |
| NEG#21 | negative | negative | 51 | female | 17.07 | 12.40 | 8.54 |
| NEG#22 | negative | negative | 45 | female | 57.86 | 37.22 | 64.01 |
| NEG#23 | negative | negative | 58 | male | 6.04 | 0.73 | 0.53 |
| NEG#24 | negative | negative | 27 | male | 30.77 | 16.88 | 35.36 |
| NEG#25 | negative | negative | 54 | male | 9.62 | 2.72 | 0.90 |
| NEG#26 | negative | negative | 58 | male | 8.16 | 2.75 | 2.59 |
| NEG#27 | negative | negative | 36 | female | 60.93 | 70.62 | 90.54 |
| NEG#28 | negative | negative | 50 | male | 18.53 | 9.34 | 5.66 |
| NEG#29 | negative | negative | 60 | male | 12.47 | 4.44 | 2.88 |
| NEG#30 | negative | negative | 67 | male | 42.86 | 67.78 | 59.50 |
| NEG#31 | negative | negative | 54 | male | 10.23 | 1.01 | 1.31 |
| NEG#32 | negative | negative | 65 | male | 24.62 | 29.67 | 23.18 |
| NEG#33 | negative | negative | 50 | male | 14.70 | 7.76 | 3.83 |
| NEG#34 | negative | negative | 36 | female | 47.34 | 43.99 | 55.12 |
| NEG#35 | negative | negative | 77 | male | 7.39 | 2.71 | 0.86 |
| NEG#36 | negative | negative | 28 | female | 49.03 | 34.90 | 50.26 |
| NEG#37 | negative | negative | 71 | female | 35.30 | 20.02 | 31.29 |
| NEG#38 | negative | negative | 39 | female | 39.79 | 55.14 | 58.86 |
| NEG#39* | negative | positive | 66 | female | 47.13 | 77.83 | 64.23 |
| NEG#40 | negative | negative | 67 | male | 0.65 | 2.93 | 0.04 |
| NEG#41 | negative | negative | 63 | male | 14.18 | 0.37 | 0.29 |
| NEG#42 | negative | negative | 39 | female | 14.25 | 4.38 | 2.72 |
| NEG#43 | negative | negative | 48 | male | 7.09 | 0.35 | 0.42 |
| NEG#44 | negative | negative | 61 | male | 13.39 | 2.87 | 6.93 |
| NEG#45 | negative | negative | 69 | male | 7.74 | 2.69 | 1.19 |
| POS#1 | positive | positive | 34 | female | 44.56 | 46.99 | 67.27 |

| | | | | | | | |
|---|---|---|---|---|---|---|---|
| POS#2 | positive | positive | 61 | female | 79.69 | 59.56 | 97.61 |
| POS#3 | positive | positive | 56 | female | 34.39 | 17.01 | 44.99 |
| POS#4 | positive | positive | 57 | female | 24.04 | 10.30 | 23.14 |
| POS#5 | positive | positive | 34 | female | 88.49 | 96.69 | 95.84 |
| POS#6 | positive | positive | 55 | male | 39.32 | 49.76 | 57.65 |
| POS#7 | positive | positive | 68 | male | 17.55 | 22.39 | 7.55 |
| POS#8 | positive | positive | 55 | male | 53.37 | 84.53 | 63.11 |
| POS#9 | positive | positive | 61 | male | 18.66 | 24.20 | 72.38 |
| POS#10* | positive | negative | 54 | male | 24.10 | 1.74 | 20.23 |
| POS#11 | positive | positive | 61 | male | 48.78 | 72.27 | 61.81 |
| POS#12 | positive | positive | 61 | female | 98.98 | 99.99 | 99.98 |
| POS#13 | positive | positive | 56 | male | 65.83 | 88.58 | 94.11 |
| POS#14 | positive | positive | 60 | male | 97.00 | 99.99 | 99.99 |
| POS#15 | positive | positive | 57 | male | 22.42 | 4.69 | 6.76 |
| POS#16 | positive | positive | 61 | male | 28.02 | 7.09 | 17.17 |
| POS#17 | positive | positive | 60 | male | 76.48 | 91.58 | 98.88 |
| POS#18* | positive | negative | 67 | male | 0.17 | 0 | 0 |
| POS#19* | positive | negative | 68 | male | 29.32 | 2.15 | 29.92 |
| POS#20* | positive | negative | 64 | male | 23.80 | 2.77 | 18.03 |
| POS#21* | positive | negative | 40 | male | 10.86 | 5.32 | 15.24 |
| POS#22* | positive | negative | 67 | male | 36.89 | 87.28 | 31.25 |
| POS#23* | positive | negative | 56 | male | 12.58 | 5.13 | 11.20 |
| POS#24* | positive | negative | 69 | male | 16.28 | 3.54 | 6.86 |
| POS#25 | positive | positive | 58 | male | 93.19 | 99.49 | 98.44 |
| POS#26* | positive | negative | 61 | male | 24.67 | 40.20 | 14.52 |
| POS#27 | positive | positive | 57 | male | 3.41 | 0.65 | 0.14 |
| POS#28 | positive | positive | 63 | male | 47.99 | 68.72 | 84.00 |
| POS#29 | positive | positive | 74 | male | 71.98 | 87.67 | 91.69 |
| POS#30 | positive | positive | 51 | female | 47.74 | 35.69 | 66.47 |
| POS#31 | positive | positive | 38 | female | 94.92 | 98.31 | 99.26 |
| POS#32* | positive | negative | 38 | female | 8.13 | 2.83 | 2.37 |
| POS#33 | positive | positive | 75 | female | 43.54 | 52.07 | 51.96 |
| POS#34 | positive | positive | 55 | male | 94.9 | 99.02 | 99.86 |
| POS#35 | positive | positive | 35 | female | 39.19 | 8.41 | 38.33 |

* represents the mismatch between cytological and histopathological results.

**Table S4 | Performance comparisons between K-PCA and ML-PCA methods.**

| Phenotyping methods | Diagnostic models | Sensitivity | Specificity | Accuracy | NPV[a] | PPV[b] |
|---|---|---|---|---|---|---|
| K-PCA | LG | 81.4% | 85.0% | 83.75% | 90% | 73.3% |
| | SVM | 77.1% | 80.4% | 78.75% | 87.5% | 65.6% |
| | LDA | 81.4% | 75.5% | 77.5% | 88.8% | 62.8% |
| ML-PCA | LG | 62.9% | 88.7% | 80% | 82.5% | 73.9% |
| | SVM | 55.5% | 81.14% | 72.5% | 78.2% | 60% |
| | LDA | 55.5% | 86.8% | 76.25% | 79.3% | 68.2% |

[a]NPV: negative predictive value, [b]PPV: positive predictive value.

**Table S5 | Comparisons between our SRC results and conventional cytology, histopathology, respectively.**

| Conventional Cytology[a, b] | Histopathology[a, b] | SRC[a, b] | Number of patients |
|---|---|---|---|
| CY+ | PM+ | SRC+ | 20 |
| CY+ | PM+ | SRC- | 5 (false negative) (see Fig. S12) |
| CY+ | PM- | SRC+ | 0 |
| CY+ | PM- | SRC- | 10 |
| CY- | PM+ | SRC+ | 2 (see Fig. S13) |
| CY- | PM+ | SRC- | 0 |
| CY- | PM- | SRC+ | 8 (false positive) |
| CY- | PM- | SRC- | 35 |

[a]+: positive, [b]-: negative.

**Table S6 | Image acquisition and processing workflow by task, software, and description.**

| | Task | Commercial (software) or custom (Github code name) | Description | Contributions / improvements |
|---|---|---|---|---|
| Imaging acquisition and stitching | Image acquisition with XY mapping | Commercial *Sciscan* 1.0 by LabVIEW | Public source code for large-area scanning imaging | Implement XY mapping with customized stage |
| | Image stiching with mapping to absolute coordinate system | Customized program by *MATLAB* | Seamless stitching of image tiles into a large FOV | Import variable overlay height and width pixel numbers when stitching large FOV |
| Single cell analysis | Cell segmentation | Commercial *Stardist* model facilitated by python TensorFlow library | Public source code for H&E or dye-staining cell segmentation | Pre-processing and post processing optimization for multicolor SRS images |
| | Feature extraction | Customized program by python *Skimage* library | Extraction of raw features of morphology and composition | Single cell features based on multiple masks |
| | Cell classification | Customized program by python *Sklearn* library | Cell phenotyping and identify significant marker cells | Unsupervised clustering using single cell-based features to get significant feature components. Meanwhile, cell phenotyping based on supervised methods |
| PM detection | PM detection of patients | Customized program by Python *Sklearn* library | Patient PM detection with feature matrix | SVM, LDA, LG models with leave-one-out cross-validation. Enable PM detection with positive probability for each patient |
| | Evaluation for the hybrid K-PCA algorithm, cell segmentation and classification | Customized program by Python *Sklearn*/ *roifile* library | Positive PM probability output, confusion matrix and ROC curves output comparing with conventional cytology and histopathology results | Compare the results from CY, PM and SRC to evaluate the improvement of our SRC method |
